# Fast Transport of RNA Granules by Direct Interactions with KIF5A/KLC1 Motors Prevents Axon Degeneration

**DOI:** 10.1101/2020.02.02.931204

**Authors:** Yusuke Fukuda, Maria F. Pazyra-Murphy, Ozge E. Tasdemir-Yilmaz, Yihang Li, Lillian Rose, Zoe C. Yeoh, Nicholas E. Vangos, Ezekiel A. Geffken, Hyuk-Soo Seo, Guillaume Adelmant, Gregory H. Bird, Loren D. Walensky, Jarrod A. Marto, Sirano Dhe-Paganon, Rosalind A. Segal

## Abstract

Complex neural circuitry requires stable connections formed by lengthy axons. To maintain these functional circuits, fast transport delivers RNAs to distal axons where they undergo local translation. However, the mechanism that enables long distance transport of non-membrane enclosed organelles such as RNA granules is not known. Here we demonstrate that a complex containing RNA and the RNA-binding protein (RBP) SFPQ interacts directly with a tetrameric kinesin containing the adaptor KLC1 and the motor KIF5A. We show that binding of SFPQ to KIF5A/KLC1 motor complex is required for axon survival and is impacted by KIF5A mutations that cause Charcot-Marie-Tooth (CMT) Disease. Moreover, therapeutic approaches that bypass the need for local translation of SFPQ-bound proteins prevent axon degeneration in CMT models. Collectively, these observations show that non-membrane enclosed organelles can move autonomously and that replacing axonally translated proteins provides a therapeutic approach to axonal degenerative disorders.

## Introduction

Sensory and motor neurons transmit signals through axons than can exceed a meter in length. Therefore, many axonal functions, including axonal survival pathways, depend on proteins that are locally translated and replenished in axon terminals. Localized protein synthesis is enabled by the initial assembly of mRNAs and RNA-binding proteins (RBPs) into ribonucleoprotein (RNP) granules which occurs within the cell soma, transport of these RNA granules to axon endings, and subsequent release of RNA for local protein synthesis (Das, Singer, & Yoon, 2019). Human mutations that disrupt RNA granule formation, interfere with cytoskeletal structures, or alter activity of intracellular motors and thus interfere with granule transport are a major cause of neurologic diseases including amyotrophic lateral sclerosis (ALS), hereditary spastic paraplegia (HSP) and Charcot-Marie-Tooth (CMT) disease. Degeneration of axons occurs early in such neurodegenerative disorders and precedes cell death of the affected neurons.

While transport of RNA granules to axons is an important step in the homeostasis of RNA and proteins in axons, the mechanism by which these non-membrane enclosed organelles are transported by microtubule-dependent motors is not yet understood. A recent study revealed one mechanism for long-range axonal transport in which *actin*-containing RNA granules “hitchhike” on lysosomes (Liao et al., 2019). However, it is not known if this represents a uniform mechanism for transport of diverse RNA-granules, or whether some types of RNA granules can be transported by motors independently of membrane containing structures.

Splicing factor proline/glutamine-rich (SFPQ) is a ubiquitous RBP that has critical functions in axons of both sensory and motor neurons (Cosker, Fenstermacher, Pazyra-Murphy, Elliott, & Segal, 2016; Pease-Raissi et al., 2017; Thomas-Jinu et al., 2017). In sensory neurons, SFPQ assembles neurotrophin-regulated transcripts, such as *bclw* and *lmnb2,* to form RNA granules, and is required for axonal localization of these mRNAs and their subsequent translation (Cosker et al., 2016; Pease-Raissi et al., 2017). Similar to many other RBPs, SFPQ contains an intrinsically disordered region (**Introduction-figure supplement 1**) and has been demonstrated to be a component of large RNA transport granules in neurons (Kanai, Dohmae, & Hirokawa, 2004). Thus, loss of SFPQ leads to depletion of axonal mRNAs and results in axon degeneration in dorsal root ganglion (DRG) sensory neurons (Cosker et al., 2016). Similarly, SFPQ is critical for the development and maintenance of motor neuron axons (Thomas-Jinu et al., 2017). Missense mutations in the coiled coil region of SFPQ have been identified that cause familial ALS and impair the localization of SFPQ within distal axon segments (Thomas-Jinu et al., 2017). However, we do not yet understand the mechanisms by which this RBP organizes mRNA transport granules that can then move rapidly to distal axons where the mRNA cargos are released and translated.

Three distinct, but closely related genes, *KIF5A, KIF5B* and *KIF5C,* encode the conventional kinesin-1 family of motor heavy chains, which are required for anterograde axonal transport of diverse organelles. Mutations in *KIF5A* cause axonal degenerative disorders including CMT Type 2D (CMT2D), HSP and ALS (Millecamps & Julien, 2013; Sleigh, Rossor, Fellows, Tosolini, & Schiavo, 2019). It has been proposed that *KIF5A* mutations may cause neurologic diseases by affecting transport efficiency overall. However, mutations in *KIF5B* or *KIF5C* do not cause similar neurologic disorders, suggesting that mutations in *KIF5A* may instead initiate disease due to impaired transport of KIF5A-specific cargo(s). Here, we show that the degeneration of axons in KIF5A or SFPQ-mutant neurons reflects failure to transport a specific non-membrane enclosed organelle rather than a general loss of transport. We identify this cargo as SFPQ-RNA granules and show that a stable small peptide that mimics the function of a locally translated protein can rescue degeneration caused by defective axonal transport.

## Results

### SFPQ granule, a non-membrane enclosed organelle, undergoes fast axonal transport

The RBP SFPQ is found in both cell bodies and axons of sensory neurons. However, the mechanisms by which SFPQ and its critical RNA cargos are transported between these two locations is not known. We utilized live cell imaging of DRG sensory neurons expressing Halo-tagged SFPQ to directly visualize transport dynamics **(Video 1)**. Fluorescent signal was enriched in the nucleus and was also evident as discrete granules in the soma and axons, a pattern similar to the distribution of endogenous SFPQ (Cosker et al., 2016). Consistent with the presence of intrinsically disordered regions within the SFPQ coding sequence, Halo-tagged SFPQ granules exhibited liquid like properties during time-lapse imaging (Gopal, Nirschl, Klinman, & Holzbaur, 2017), as the size and shape of SFPQ granules remained constant at approximately 1 μm in diameter during the stationary phase, but the granules expanded and elongated as they move (**Figure 1A and 1B**). The majority of the Halo-tagged SFPQ granules in axons were motile, either moving by retrograde transport (∼48%), or anterograde transport (∼28%), with the remainder in stationary phase (∼25%) (**Figure 1C and Figure 1-figure supplement 1A**). SFPQ granules exhibit an average anterograde velocity of 0.89 ± 0.08 μm/sec and average anterograde cumulative displacement of 21.02 ± 2.49 μm, with an average retrograde velocity of 0.80 ± 0.04 μm/sec retrograde and average retrograde cumulative displacement of 32.02 ± 2.45 μm (**Figure 1D, Figure 1-figure supplement 1B-E**). Together, the velocity and the characteristics of movement indicate that the SFPQ-granules are non-membrane enclosed organelles that move in both directions by microtubule-dependent fast axonal transport, using a kinesin motor for anterograde and the more highly processive dynein motor for retrograde movements.

**Figure 1.**
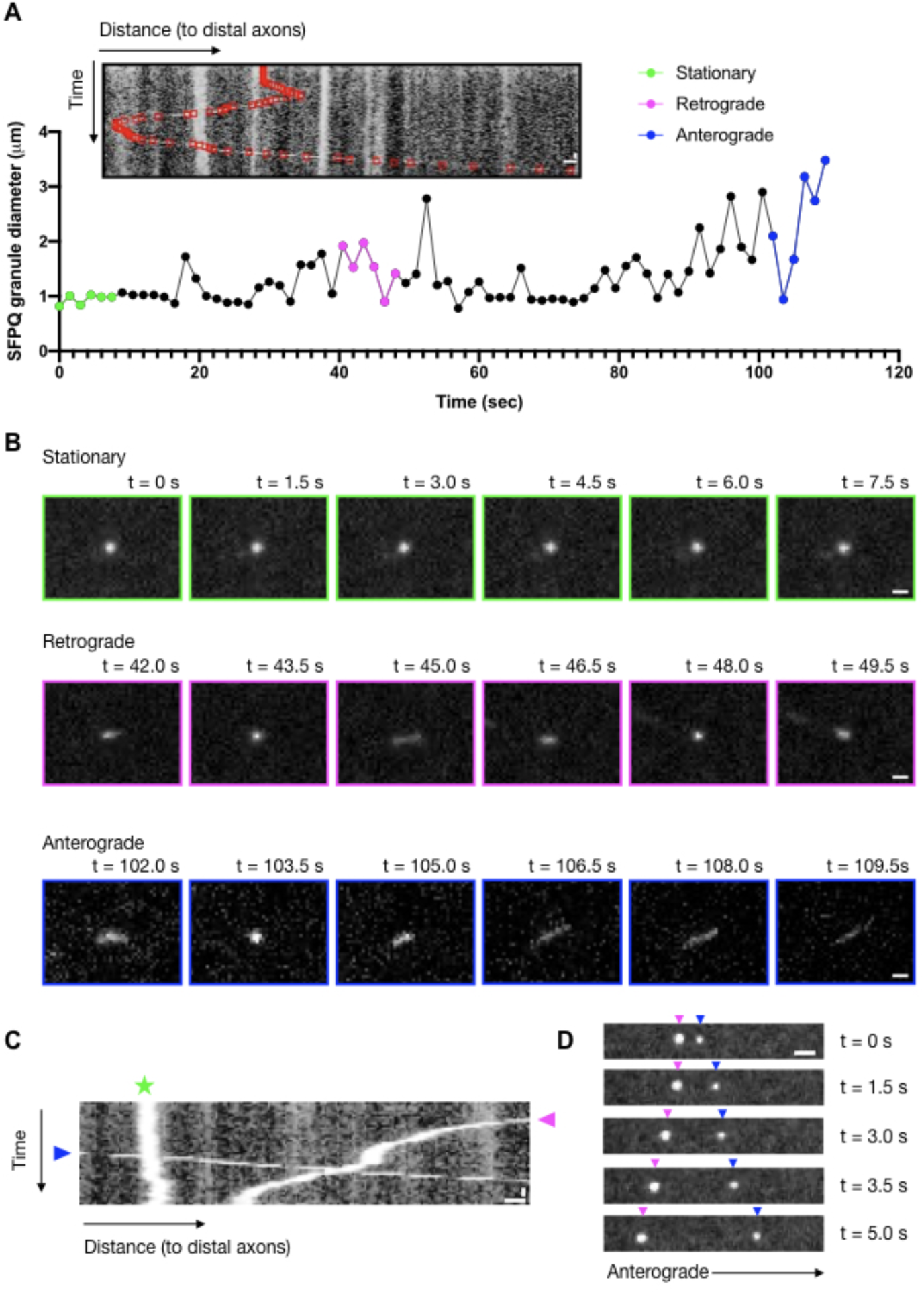
SFPQ granule, a non-membrane enclosed organelle, undergoes fast axonal transport. (A) Kymograph of a Halo-SFPQ granule transitioning through stationary, retrograde and anterograde transport and its diameter plotted over time. Scale bars: 2 μm and 6 sec. (B) Time-lapse images of the corresponding Halo-SFPQ granule from (A) in stationary (green), retrograde (magenta) and anterograde (blue) phase. Scale bars: 1 μm. (C) Kymograph depicting representative Halo-SFPQ in stationary (green star), anterograde (blue arrowhead) and retrograde (magenta arrowhead) phase. Scale bars: 2 μm and 6 sec. (D) Representative time-lapse images of Halo-SFPQ in axons of DRG sensory neurons moving in anterograde (blue arrowhead) or retrograde (magenta arrowhead) direction. Scale bar: 2 μm.

### SFPQ preferentially binds to KIF5A/KLC1 motor complex

The anterograde kinesins involved in axon transport are formed by two dimers of kinesin heavy chain (KHC or KIF5) and two dimers of kinesin light chain (KLC) (**Figure 2A**). The KIF5 family is encoded by three distinct genes, *KIF5A*, *KIF5B* and *KIF5C,* and the genome also contains several light chains, KLC1-4. To identify motors that associate with SFPQ and might enable transport of these RNA granules, we took an unbiased approach in which we immunoprecipitated endogenous SFPQ from DRG neurons and used mass spectrometry to analyze the co-precipitated components. We detected known interactors, including the *Drosophila* behavior/human splicing (DBHS) protein family members NONO and PSPC1 in the precipitated proteins (**Supplementary File 1**). We then analyzed the relative abundance of KIF5A, KIF5B and KIF5C (**Figure 2B**) and KLC1, KLC2 (**Figure 2C**) in the SFPQ immunoprecipitates using a targeted mass spectrometry approach. We found that KIF5A and KLC1 were each highly enriched over the other proteins (KIF5B or KIF5C and KLC2, respectively) as measured across three independent experiments (**Figure 2B, 2C and Figure 2-figure supplement 1A**). In contrast, when we purified endogenous KLC1 from DRG neurons and analyzed the composition of the resulting immunoprecipitate by mass spectrometry, we observed that all three KIF5 proteins were present at approximately the same abundance in KLC1 immunoprecipitate (**Figure 2-figure supplement 1A and 1B**). Similarly, validated antibodies specific to each of the three KIF5s (**Figure 2-figure supplement 1C and 1D**) corroborate that SFPQ preferentially co-immunoprecipitates with KIF5A, rather than the closely related KIF5B or KIF5C (**Figure 2D and 2E**), and with KLC1 rather than KLC2 (**Figure 2F**). Together these results suggest the possibility that KIF5A/KLC1 tetramers may be the distinctive motors responsible for rapid anterograde transport of SFPQ-RNA granules.

**Figure 2.**
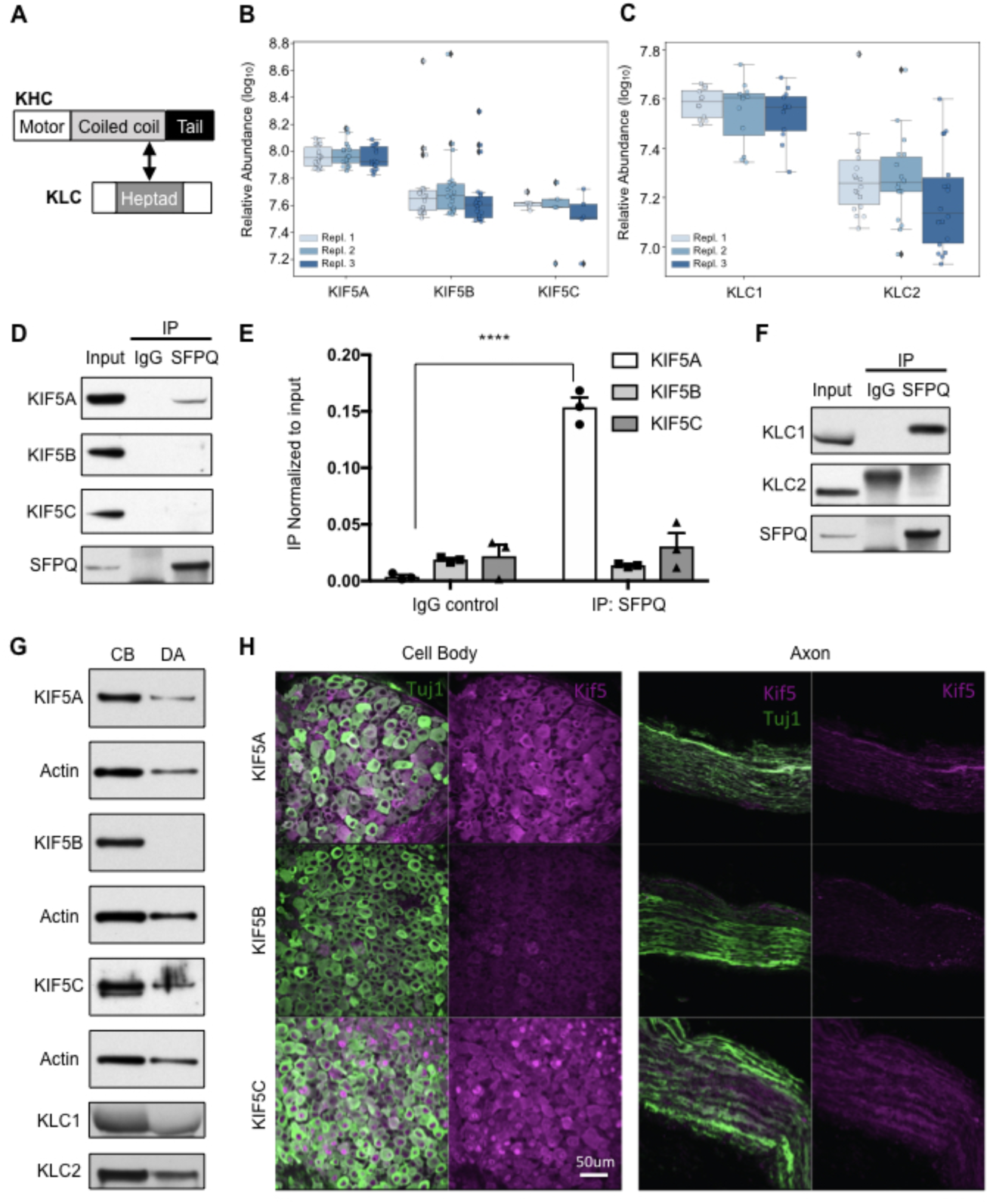
SFPQ preferentially binds to KIF5A/KLC1 motor complex. (A) Schematic of domains of kinesin heavy chain (KHC) and kinesin light chain (KLC) and the interacting region between the heptad repeat of KLC and the coiled coil of KHC. (B and C) Box and whisker plot showing the relative abundance of KIF5A, KIF5B and KIF5C (B); and KLC1 and KLC2 (C) peptides derived from parallel reaction monitoring mass spectrometry. Data were acquired across three independent SFPQ immunoprecipitations (IPs). (D) IP of endogenous SFPQ from DRG sensory neuron protein lysate and blotted against endogenous KIF5A, KIF5B and KIF5C; IgG serves as control IP. (E) Quantification of pull down in (D) relative to input; ****p < 0.0001 by one way ANOVA; n = 3 independent IPs; data represent mean ± s.e.m. (F) IP of endogenous SFPQ from DRG sensory neuron protein lysate and blotted against endogenous KLC1 and KLC2; IgG serves as control IP. (G) Western blot of DRG neuron lysates of cell body (CB) and distal axons (DA) prepared from compartmented Campenot cultures probed against endogenous KIF5A, KIF5B, KIF5C, KLC1 and KLC2; actin serves as loading control. (H) Representative staining of endogenous KIF5A, KIF5B and KIF5C in DRGs and sciatic nerve of P1 mice; n = 4 independent staining of tissues; scale bar 50 μm; Tuj1 (Green), KIF5 (Magenta).

A previous study of kinesin-1 motors demonstrates that overexpressed KIF5A, B and C can all traffic to axons, but KIF5A is excluded from dendrites (Lipka, Kapitein, Jaworski, & Hoogenraad, 2016). DRG sensory neurons are pseudo-unipolar in morphology and so have no dendrites, but instead consist of a cell body with a T-shaped axon. To determine whether the intracellular distribution of kinesins is consistent with the hypothesis that KIF5A/KLC1 is the motor that transports SFPQ-RNA granules, we cultured DRG sensory neurons in compartmented cultures and collected protein lysates distinctly from cell body (CB) and distal axon (DA) compartments. While KIF5B preferentially localizes in the CB compartment, KIF5A and KIF5C localize to both CB and DA compartments, as do KLC1 and KLC2 (**Figure 2G**). Immunostaining of KIF5s in DRG neurons cultured in microfluidic chambers as well as DRGs and sciatic nerves *in vivo* displayed a similar localization pattern, with KIF5B largely excluded from axons (**Figure 2H and Figure 2-figure supplement 2**). Together these data demonstrate that KIF5A and KLC1 are appropriately localized to mediate transport of SFPQ-RNA granules from cell bodies to distal axons, and thus KIF5A/KLC1 may represent a specialized motor for these non-membrane enclosed organelles.

### RNase prevents SFPQ-RNA binding to KIF5A/KLC1

As RNA is a critical component of the large SFPQ-containing granules that move rapidly within the axons, we asked whether SFPQ that binds to KIF5A/KLC1 is also associated with RNA cargos. We expressed HA-SFPQ, FLAG-KIF5A and KLC1-Myc in HEK293T cells, and treated the cell lysates with RNase, or vehicle control, and then immunoprecipitated for HA-SFPQ. Strikingly we find that RNase treatment impeded the interaction between SFPQ and KIF5A-KLC1 (**Figure 3A-C**), demonstrating that SFPQ only binds KIF5A/KLC1 when it is associated with RNA. These findings suggest that KIF5A/KLC1 bind and transport SFPQ when it is part of a large RNP transport granule.

**Figure 3.**
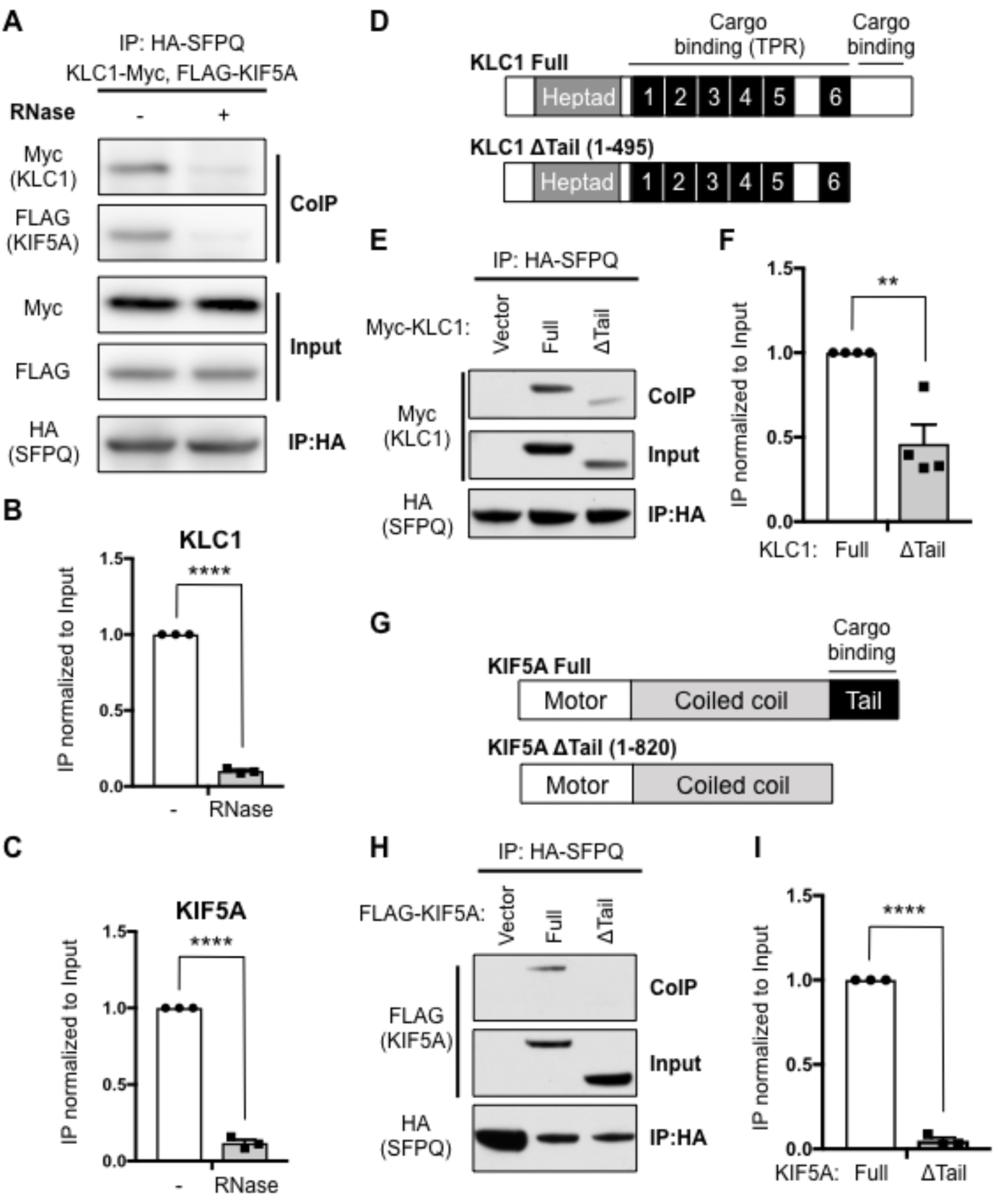
RNase prevents SFPQ-RNA binding to KIF5A/KLC1. (A) HEK 293T cells transfected with HA-SFPQ, FLAG-KIF5A and KLC1-Myc and lysates were treated with or without RNase. HA was IPed and blotted for HA, Myc and FLAG. (B) Quantification of pull down of KLC1-Myc in (A) relative to input; ****p < 0.0001 by unpaired two-tailed t test; n = 3; data represent mean ± s.e.m. (C) Quantification of pull down of FLAG-KIF5A in (A) relative to input; ****p < 0.0001 by unpaired two-tailed t test; n = 3; data represent mean ± s.e.m. (D) Schematic of the indicated constructs for KLC1; Heptad, Heptad repeat; TPR, tetratricopeptide repeat. (E) HEK 293T cells transfected with HA-SFPQ with empty vector, full length WT or tail-truncated Myc-KLC1. HA was IPed and blotted for Myc and HA. (F) Quantification of pull down in (E) relative to input; **p = 0.0033 by unpaired two-tailed t test; n = 4; data represent mean ± s.e.m. (G) Schematic of the indicated constructs for KIF5A. (H) HEK 293T cell transfected with HA-SFPQ with empty vector, full length WT or tail-truncated FLAG-tagged KIF5A. HA was IPed and blotted for FLAG and HA. (I) Quantification of pull down in (H) relative to input; ****p < 0.0001 by unpaired two-tailed t test; n = 3; data represent mean ± s.e.m.

Based on the above findings, we hypothesize that KIF5A/KLC1 distinctively mediates fast transport of critical SFPQ-RNA granules from the cell soma to the axons. To identify the structural basis for this specificity, we first asked whether the highly divergent C-terminal tail regions of KLC1 and KIF5A are required for SFPQ binding. When we overexpressed either Myc-tagged WT KLC1 or its C-terminal mutant (ΔTail) in HEK 293T cells, and assessed binding to SFPQ by co-precipitation studies, we find that the binding is reduced by approximately 50% in the absence of the C-terminal region of KLC1 (**Figure 3D-F**). Similarly, truncation of the highly variable tail region of KIF5A nearly abolished its interaction with SFPQ (**Figure 3G-I**) although this did not prevent binding of KIF5A to KLC1 (**Figure 3-figure supplement 1**). Together these data demonstrate that SFPQ binding to KIF5A/KLC1 is enabled by the highly divergent C-terminal regions of both KLC1 and KIF5A.

### SFPQ directly binds to KLC1 through a Y-acidic motif within its coiled coil domain

Evidence that SFPQ binding to KIF5A/KLC1 requires concurrent binding to RNA, suggests that this RBP may directly link RNA transport granules to kinesin motors, enabling these RNA granules to move autonomously without a membranous platform. To develop novel tools for testing this possibility, we sought to identify specific mutations in SFPQ that impair binding to kinesin. SFPQ contains a Y-acidic motif that is evolutionary conserved and closely matches the Y-acidic sequence of JIP1 that connects membranous organelles to KLC1 (Nguyen et al., 2018; Pernigo et al., 2018) (**Figure 4A and Figure 4-figure supplement 1**). When we mutated the critical tyrosine residue within the motif to alanine (Y527A) the binding between SFPQ and KIF5A/KLC1 was dramatically reduced, demonstrating that this Y-acidic motif is required for binding to KIF5A/KLC1 motor complex (**Figure 4B and 4C**). To assess whether SFPQ binds directly to KLC1 without requiring a membranous organelle or an adaptor component, we purified human KLC1 and used isothermal titration calorimetry (ITC) to test direct binding by a long SFPQ peptide that spans the Y-acidic motif. The SFPQ peptide binds directly to KLC1 with a Kd of 3.8 ± 2.3 μM and a binding stoichiometry of 1 (**Figure 4D and Table 1**). Consistent with data from co-immunoprecipitation studies above, Y527A mutation prevents the SFPQ peptide from binding directly to KLC1 in ITC assays (**Figure 4D**). Thermodynamic parameters of ITC measurements are summarized in Table 1. Together these data demonstrate that SFPQ directly binds to KLC1 in a process that relies on the Y-acidic motif and is abrogated by the Y527A mutation. These studies suggest that SFPQ RNA-transport granules might directly associate with microtubule-dependent motors rather than requiring a membrane platform for intracellular transport.

**Figure 4.**
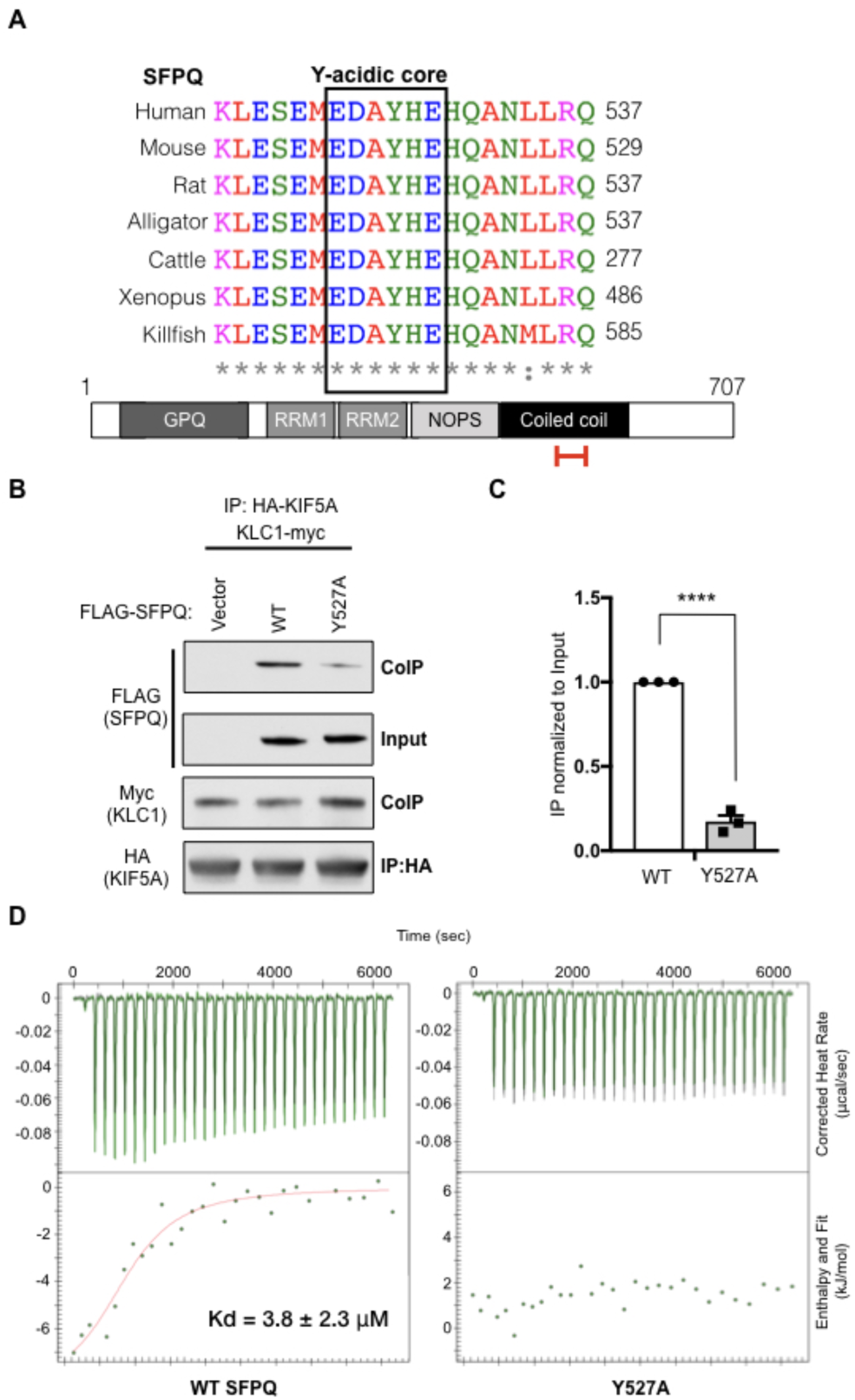
SFPQ directly binds to KLC1 through a Y-acidic motif within its coiled coil domain. (A) Alignment of the sequence within the coiled coil domain of SFPQ containing the Y-acidic motif. On the bottom; schematic of the domains of SFPQ. Red bracket indicates the region containing the Y-acidic motif. GPQ, glycine proline glutamine-rich; RRM, RNA recognition motif; NOPS, NONA/paraspeckle domain. (B) HEK 293T cells transfected with HA-KIF5A, KLC1-Myc and with either empty vector, full length WT or Y527A FLAG-tagged SFPQ. HA was IPed and blotted for FLAG, Myc and HA. (C) Quantification of pull down in (B) relative to input; ****p < 0.0001 by unpaired two-tailed t test; n = 3; data represent mean ± s.e.m. (D) Isothermal titration calorimetry (ITC) measurements of the reference KLC1 (TPR1-6) fragment with either the WT peptide (ESEMEDA<u>Y</u>HEHQANLLR) or the Y-acidic mutant, Y527A, peptide (ESEMEDA<u>A</u>HEHQANLLR) of SFPQ.

**Table 1.** ITC parameters between KLC1 TPR1-6 fragment and WT SFPQ or Y527A Y-acidic mutant peptide.

| **KLC1**<br>**TPR1-6**<br>**(Cell)** | **SFPQ**<br>**(Syringe)** | **N** | **Kd**<br>**(μM)** | **ΔH**<br>**(kJ/mol)** | **ΔS**<br>**(J/mol·K)** |
| --- | --- | --- | --- | --- | --- |
| | **WT** | 0.964 ± 0.143 | 3.833 ± 2.310 | -8.580 ± 1.834 | 74.92 |
|  | **Y527A** | No Binding |  |  |  |

### Direct binding of SFPQ to KIF5A/KLC1 is required for its transport in axons

The Y527A mutant of SFPQ provides a tool that can be used to ask whether the direct interaction between SFPQ and kinesin motors is responsible for autonomous transport of SFPQ-RNA granules along microtubules. Such a transport mechanism would contrast with previous models for transport of non-membrane enclosed granules, as RNA granules that contain β-actin *mRNA* “hitchhike” on membrane platforms, and do not move autonomously (Liao et al., 2019). To determine whether the direct binding of SFPQ-RNA granules to kinesin mediates fast axonal transport, we expressed Halo-tagged WT or Y527A SFPQ within DRG neurons grown in microfluidic chambers. In neurons expressing the Y527A mutant **(Video 2)**, the number of SFPQ particles localized to distal axons was reduced by ∼50%, suggesting that direct binding of SFPQ to KIF5A/KLC1 is required for the redistribution of SFPQ-granules from the cell bodies to distal axons (**Figure 5A and 5B**). Moreover, among the SFPQ granules that reached the axons, the Y527A mutant SFPQ exhibited a ∼50% reduction in the percentage of time spent in anterograde axonal transport compared to WT (**Figure 5C**). Since SFPQ forms a dimer, residual movement of Y527A may reflect binding of KIF5A/KLC1 motors to dimeric SFPQ containing both endogenous WT SFPQ and the fluorescent mutant isoform (Hewage, Caria, & Lee, 2019). Together these data demonstrate that defects in the direct binding of Y527A to KIF5A/KLC1 motor complex interrupts anterograde transport of RNA granules in axons of DRG sensory neurons, suggesting that these RNA granules move autonomously and do not require a membrane platform.

**Figure 5.**
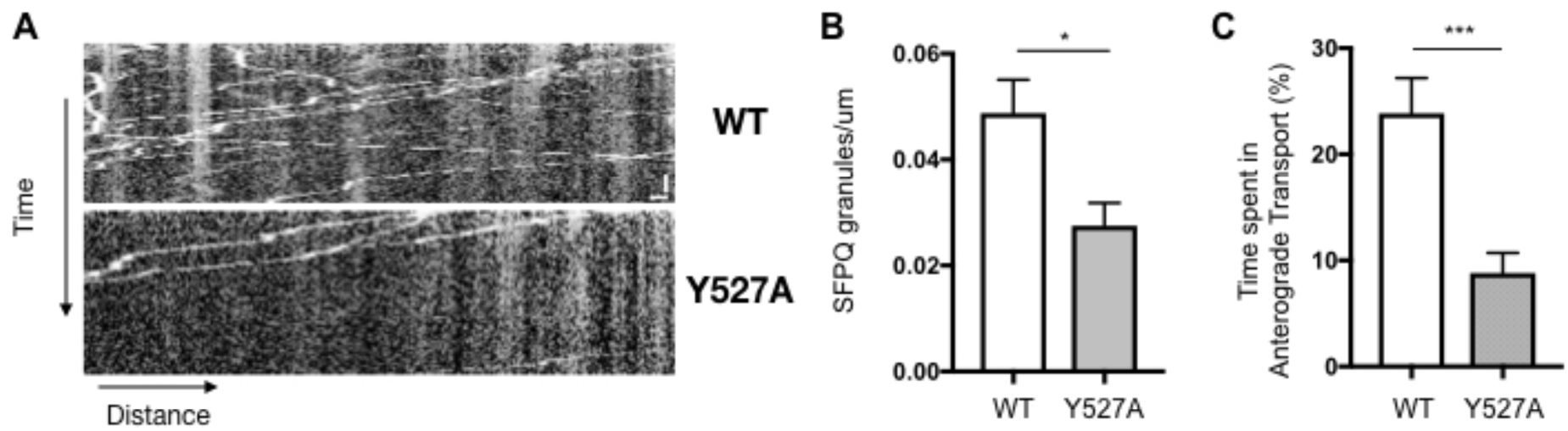
Direct binding of SFPQ to KIF5A/KLC1 is required for its transport in axons. (A) Representative kymograph of WT and Y527A Halo-tagged SFPQ. Scale Bars: 2 μm and 15 sec. (B) Average number of Halo-tagged WT and Y527A per micron of axon length. Analyzed from n = 25-33 axons from two independent experiments; *p = 0.0117; data represent mean ± s.e.m. (C) Average percentage of time spent in anterograde transport for Halo-tagged WT and Y527A in axons of DRG sensory neurons. Analyzed from n = 25-33 axons from 2 independent experiments; ***p = 0.0005; data represent mean ± s.e.m.

It is striking that the Y-acidic motifs in JIP1 and other proteins that link kinesins to membranous organelles are usually located within highly accessible regions such as the carboxy terminus, while the Y-acidic motif in SFPQ is instead located within the highly structured coiled-coil domain. These structural differences suggest that SFPQ-RNA granules may interact with kinesin in a different manner than do membranous organelles. Previous studies identified sequences within KLC1 that are not present in KLC2 and that specify binding to JIP1. One key residue in KLC1 is N343 within the TPR4 region of KLC1; mutation of this residue to a serine, as observed in KLC2, abrogates interaction between JIP1 and KLC1 (Zhu et al., 2012) (**Figure 5-figure supplement 1A**). To determine whether N343 on KLC1 also mediates binding to SFPQ, we expressed myc-tagged WT or N343S KLC1 together with HA-tagged SFPQ in HEK 293T cells. We find that KLC1-N343S did not alter binding to SFPQ, in clear contrast to JIP1 (**Figure 5-figure supplement 1B**), suggesting that the way in which SFPQ-RNA granules bind to kinesin motors for transport differs from that observed with JIP1.

### Defect in KIF5A-driven transport of SFPQ leads to axon degeneration in DRG sensory neurons

To determine the physiologic consequences of impeding SFPQ transport by KIF5A/KLC1, we leveraged the SFPQ Y527A mutant (**Figure 6A and 5**). As shown previously, knockdown of SFPQ results in axon degeneration. We then assessed the ability of constructs encoding either the WT SFPQ or SFPQ-Y527A to reverse the degeneration caused by knockdown, using constructs resistant to shRNA knockdown (**Figure 6B and Figure 6-figure supplement 1**). Strikingly, expression of the WT version rescued axon degeneration caused by knockdown of SFPQ, whereas expression of SFPQ Y527A was unable to do so, and instead led to the same degree of axon degeneration observed following knockdown of SFPQ (**Figure 6C and 6D**). Taken together these data indicate that autonomous transport of SFPQ-RNA granules is required for axon survival and demonstrate that defects in kinesin-driven transport of SFPQ causes degeneration of sensory axons.

**Figure 6.**
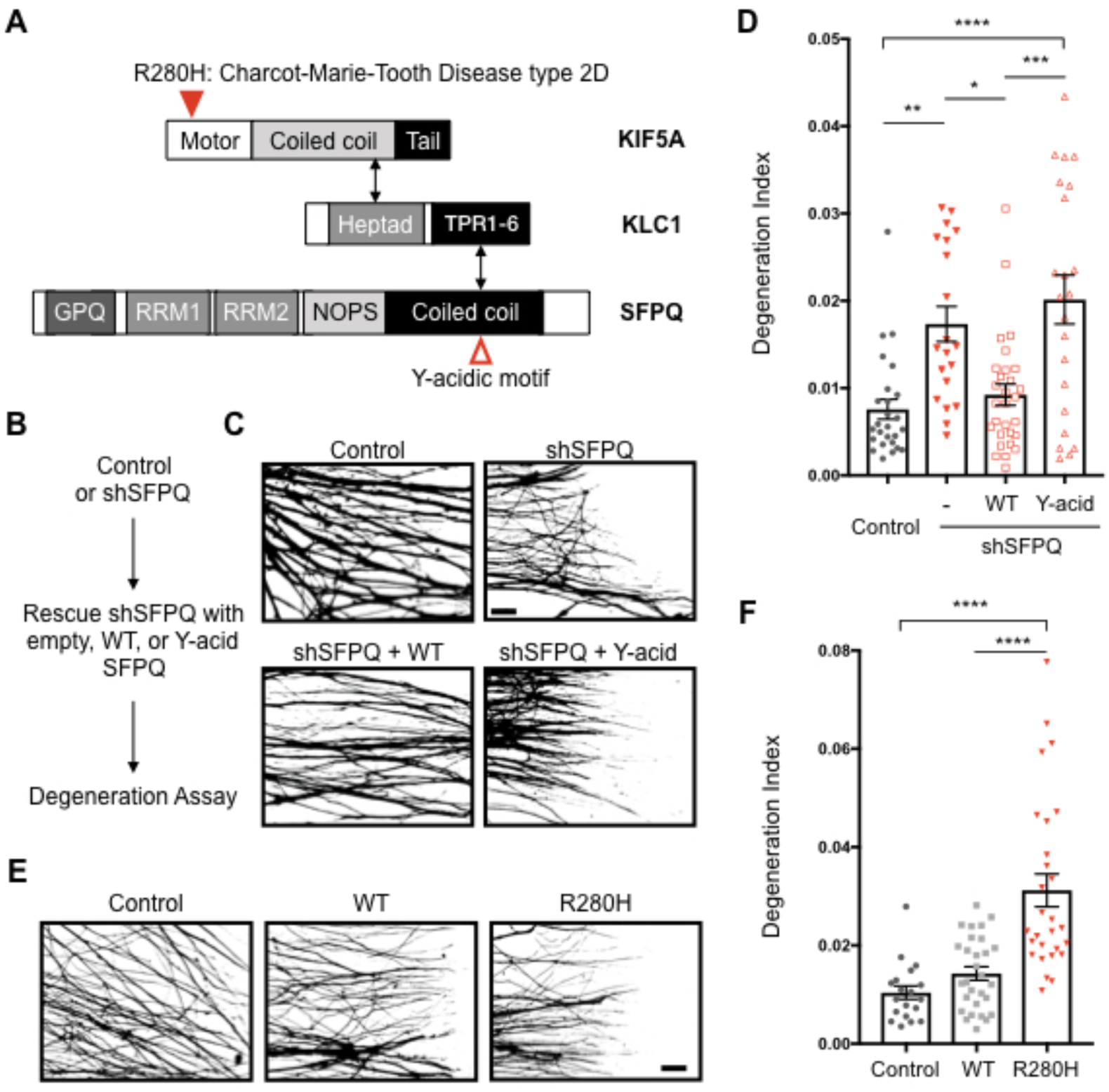
Defect in KIF5A-driven transport of SFPQ leads to axon degeneration in DRG sensory neurons. (A) A schematic representation of KIF5A, KLC1 and SFPQ. Red closed arrowhead indicates the location of R280H CMT2D mutation; and open arrowhead indicates the location of Y-acidic motif of SFPQ. (B) A flowchart of rescue experiment of degeneration assay using WT or the Y-acidic mutant of SFPQ. (C) Representative binarized Tuj1-labeled axons in compartmented cultures expressing control (n = 26) or shSFPQ rescued with empty vector (n = 20), WT (n = 29) or the Y-acidic mutant (n = 22) of SFPQ. From 3 independent experiments; Scale bar 100 μm. (D) Quantification of axon degeneration index of (C); *p = 0.0112, **p = 0.0019, ***p = 0.0002, ****p < 0.0001 by one way ANOVA; data represent mean ± s.e.m. (E) Representative binarized Tuj1-labeled axons in compartmented cultures expressing control (n = 19), WT (n = 29) or R280H (n = 28) mutant of KIF5A. From 3 independent experiments; Scale bar 100 μm. (F) Quantification of axon degeneration index of (E); ****p < 0.0001 by one way ANOVA; data represent mean ± s.e.m.

Our data indicate that KIF5A/KLC1 functions as the motor transporting SFPQ-RNA granules within DRG sensory neurons, and that this distinct transport process is required for axon survival. Thus, CMT2D mutations of KIF5A may cause defects in this specialized transport pathway and so compromise axonal health. Mutations causing HSP and CMT2D are located primarily within the motor domain of KIF5A; each distinct mutation can give rise to HSP or CMT2D or a combination of both syndromes. Two independent studies have reported a R280H mutation that results in a pure classical form of CMT2D (**Figure 6A**) (Liu et al., 2014; Nam, Yoo, Choi, Choi, & Chung, 2018). Both structural and biochemical characterization indicates that disease mutations altering this residue reduce the microtubule binding affinity, and therefore decrease transport (Dutta, Diehl, Onuchic, & Jana, 2018; Ebbing et al., 2008; Fuger et al., 2012; Jennings et al., 2017). Interestingly, despite the mutation residing within the motor domain, R280H mutation also reduced binding to SFPQ approximately 25% compared to WT KIF5A (**Figure 6-figure supplement 2A and 2B**). As CMT2D mutation within R280H not only reduces microtubule binding affinity and transport but also impacts binding to SFPQ, this pathologic mutation is likely to disproportionately affect transport of the RNA cargo transported together with SFPQ.

To model axon degeneration induced by KIF5A mutation in R280H *in vitro*, we expressed the KIF5A R280H mutation by lentivirus in DRG sensory neurons that also express the endogenous WT KIF5A. As observed in patients heterozygous for CMT2D mutations, overexpression of R280H led to axon degeneration in DRG sensory neurons (**Figure 6E and 6F**), while control studies indicate that overexpression of WT does not compromise axon integrity (**Figure 6E and 6F**). This model provides a platform for investigating the molecular changes that cause degeneration in CMT disease.

### Axon degeneration caused by CMT2D R280H KIF5A mutation can be rescued by a Bclw mimetic peptide

KIF5A motors are involved in transport of mitochondria, vesicles, and other membrane enclosed organelles as well as RNA granules; thus degeneration of axons caused by R280H KIF5A mutation could be a consequence of defect in transport of any or all of these cargos. A similar pattern of axon degeneration is observed in sensory neurons expressing the SFPQ Y527A mutant, suggesting that SFPQ-RNA granules may represent a critical cargo impacted in CMT patients with KIF5A R280H mutations. Local translation of mRNAs bound to SFPQ is a critical step that promotes axon survival; *bclw* is one such mRNA that is bound by SFPQ and is translated in axons of DRG sensory neurons (**Figure 7A**). We asked whether restoring functions downstream of the SFPQ pathway can rescue degeneration caused by KIF5A R280H mutant. Intriguingly, a BH4 peptide mimetic of Bclw introduced into axons of R280H KIF5A mutant can prevent axon degeneration in this genetic model of CMT with axonal survival returning to the levels observed in control or WT KIF5A overexpressing DRG neurons (**Figure 7B**). This effect is specific to Bclw since introducing a peptide mimetic of related Bcl2 proteins did not prevent degeneration, and degeneration remained at the level observed with the R280H mutant alone (**Figure 7C**). Taken together, these data suggest that defective transport of SFP-RNA granules containing *bclw* mRNA represents a key mechanism that underlies CMT2D-causing mutations of KIF5A, and so causes axon degeneration.

**Figure 7.**
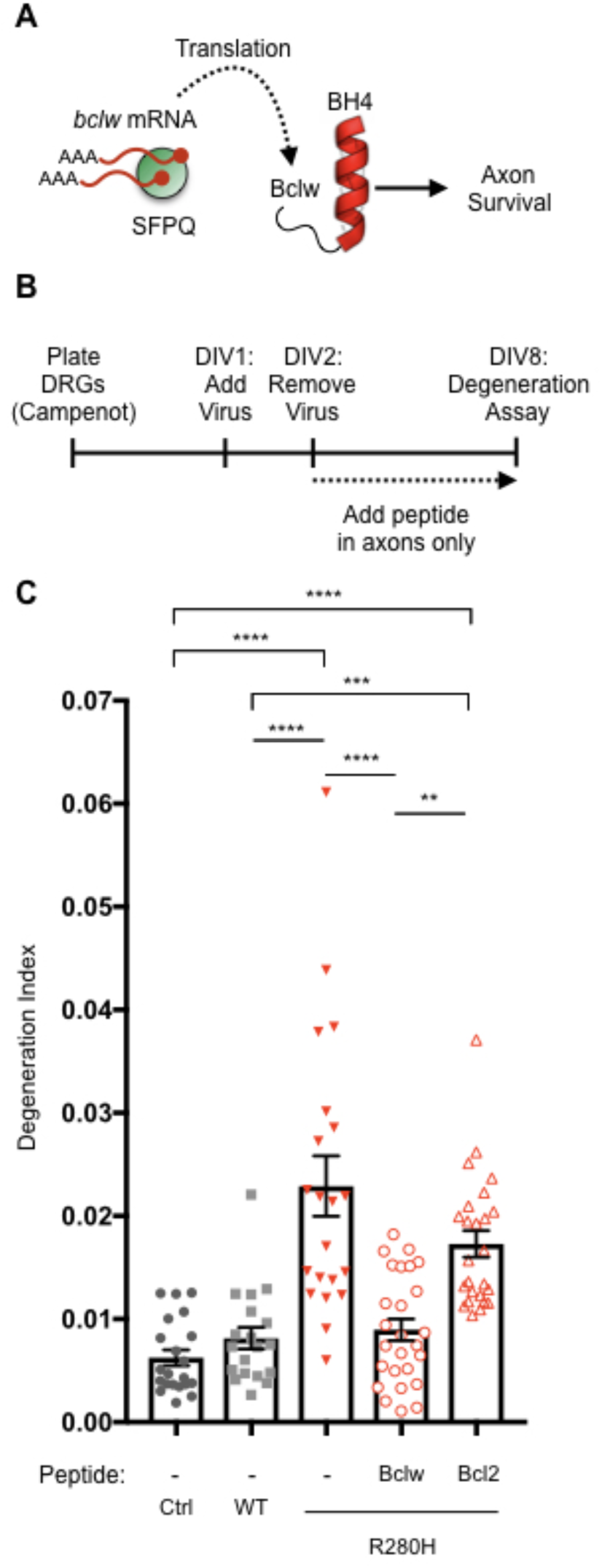
Axon degeneration caused by CMT2D R280H KIF5A mutation can be rescued by a Bclw mimetic peptide. (A) A schematic of pathway for axon survival mediated by SFPQ. (B) Flowchart of Bclw peptide rescue experiment in DRG neurons cultured in compartmented Campenot chambers. (C) Quantification of axon degeneration index of control (Ctrl; n = 21), WT KIF5A (n = 19), R280H with either no peptide (n = 21), Bclw (n = 26), or Bcl2 peptide (n = 25). From 3 independent experiments; **p = 0.0011, ***p = 0.0010, ****p < 0.0001 by one way ANOVA; data represent mean ± s.e.m.

## Discussion

Kinesins are a large family of microtubule-dependent motors that play a pivotal role in rapid intracellular transport. These motors are particularly critical in neurons that form extensive axonal and dendritic projections and so rely on fast transport across long distances. While mutations in kinesin motors cause neurodegenerative diseases affecting primary sensory and motor neurons, it is not known whether this reflects a requirement for specific kinesins in mediating transport of particular cargos within the lengthy axons that extend to peripheral targets, or whether degeneration reflects a more global loss of axonal transport. Here we show that the non-membrane enclosed RNA granules containing the RNA binding protein SFPQ selectively and directly interact with kinesin containing the KIF5A heavy chain and the cargo adaptor KLC1. Therefore, mutations in KIF5A that cause sensory neuropathy preferentially impact motility of SFPQ granules that transport *bclw* and other mRNAs, and the degeneration caused by KIF5A mutations can be prevented by a peptide that mimics the function of the locally translated protein Bclw.

Although it is widely accepted that RNA-granules are non-membrane enclosed organelles that move by microtubule-dependent transport, how such RNA-granules associate with motors and move through the axoplasm is not yet known. Our data demonstrate a direct interaction between SFPQ and the kinesin-1 cargo adaptor complex KLC1, and we show that this interaction is required for autonomous axonal transport. In contrast to this direct transport system, a recent study by Liao *et al*. demonstrated that RNA-granules containing *β-actin* mRNA rely on annexin A11 adaptor protein to hitchhike on lysosomes and thereby regulate growth cone morphology (Liao et al., 2019). As SFPQ does not bind *β-actin* but instead binds mRNAs that promote axonal survival, our findings indicate that distinct pools of RNA-granules containing different mRNAs rely on divergent modes of axonal transport.

Data from quantitative mass spectrometry and isothermal titration calorimetry demonstrate that SFPQ preferentially binds KLC1/KIF5A in DRG sensory neurons and that an evolutionarily conserved Y-acidic motif is sufficient for binding directly to KLC1. Interestingly, binding of the protein cargo/adaptors JIP1, TorsinA and SH2D6 to KLC1 require Y-acidic motifs within an unstructured part of the molecule (Nguyen et al., 2018; Pernigo et al., 2018), whereas the motif within SFPQ resides in the highly structured coiled-coil region of SFPQ (**Figure 4A and Figure 4-figure supplement 1**). As RNA interactions enable SFPQ to bind KIF5A/KLC1, formation of a RNP complex may expose the Y-acidic motif within the coiled-coil domain. Thus, the structural basis for the interaction of KIF5A/KLC1 with SFPQ-RNA granules is likely to differ from the mechanism for binding of KIF5A/KLC1 to JIP1. Consistent with this distinction, the N343S mutation of KLC1 that disrupts interaction with JIP1 (Pernigo et al., 2018; Zhu et al., 2012) has no effect on binding with SFPQ (**Figure 5-figure supplement 1B**). Instead the highly divergent C-terminal region of KLC1 affects binding to SFPQ (**Figure 3D-F**). As the carboxy terminal region of KLC1 diverges among several splice variants of KLC1 (McCart, Mahony, & Rothnagel, 2003), individual isoforms of KLC1 that are present in DRG sensory neurons may specify binding to SFPQ-RNA granules.

Similar to other phase separating RBPs, SFPQ contains an intrinsically disordered region in its N-terminal region (**Introduction-figure supplement 1**) and SFPQ assembles into large RNA transport granules, which are non-membrane enclosed organelles (Kanai et al., 2004). In the nucleus, SFPQ interacts with members of the DBHS proteins, NONO and PSPC1, and also binds RNA fragments of NEAT1_2 in order to form paraspeckle nuclear bodies, another phase separating structure (Yamazaki et al., 2018). As is the case for paraspeckle formation, RNA is a critical component of transport granules; RNase treatment leads to loss of integrity of these axonal granules (Knowles et al., 1996), and binding between SFPQ and KIF5A/KLC1 is highly sensitive to RNase treatment (**Figure 3A-C**). Binding to mRNAs such as *bclw* may induce a conformational change in SFPQ that facilitates oligomerization and phase separation and exposes the Y-acidic motif within the coiled coil domain so it can bind KLC1/KIF5A. Further structural studies investigating how specific RNA cargos regulate the conformation and binding of SFPQ to KLC1 and KIF5A will be necessary to fully evaluate this hypothesis.

Among the three *KIF5* genes encoding the kinesin-1 family of motors, the *KIF5A* gene is the only one associated with the human neurological diseases CMT2D, HSP and ALS. Our results demonstrate that binding of SFPQ to KLC1 complexed with KIF5A rather than KIF5B or KIF5C enables transport of SFPQ-RNA granules and promotes axon survival. Based on these findings, we postulate that defective transport of SFPQ-RNA granules is a major contributor to KIF5A-associated neurodegenerative disorders, rather than axon degeneration in these disorders being the result of a generalized impairment of transport.

SFPQ RNA-granules are critical for development and maintenance of axons in motor neurons as well as in sensory neurons; and human mutations in *SFPQ* and *KIF5A* have been implicated in ALS (Thomas-Jinu et al., 2017). Interestingly, the Y527 residue of the Y-acidic motif lies adjacent to the two identified ALS mutations: N533H and L534I; these two mutations lead to defects in the axonal functions of SFPQ in motor neurons (Thomas-Jinu et al., 2017) and our data suggest that this may reflect altered binding of SFPQ to KIF5A/KLC1 and altered autonomous transport of these non-membrane enclosed granules. Moreover, ALS-associated mutations in *KIF5A* transform the C-terminal region of the protein (Nicolas et al., 2018), the domain required for SFPQ binding. Therefore, altered binding and axonal transport of SFPQ may explain the axon degeneration of motor neurons in patients harboring ALS-associated mutations in *SFPQ* or *KIF5A*. A recent study by Luisier *et al*. demonstrated that aberrant localization of SFPQ is also a molecular hallmark of multiple familial and sporadic models of ALS that do not exhibit mutations in *SFPQ* or *KIF5A* (Luisier et al., 2018), suggesting that disrupted transport and axonal function of SFPQ may contribute to additional neurodegenerative disorders. Here we demonstrated that a Bclw peptide mimetic rescues axon degeneration caused by R280H KIF5A mutations (**Figure 7**), extending previous evidence that this therapeutic approach prevents paclitaxel-induced axon degeneration (Pease-Raissi et al., 2017). Taken together, our data suggests that Bclw peptide mimetics should be explored as a potential therapeutic intervention for preventing axon degeneration in multiple neurological diseases that share defects in RNA transport as part of their pathophysiology.

## Acknowledgments

We thank Thomas Schwarz and Himanish Basu (Boston Children’s Hospital, MA) for Kymolyzer software. We thank Gary Banker (Oregon Health and Science University, OR) and Marvin Bentley (Rensselaer Polytechnic Institute, NY) for KIF5A, KIF5B, KIF5C and KLC1 constructs; and Corinne Houart (Kings’s College London, United Kingdom) for SFPQ construct. We thank the Segal lab, Charles Stiles and Michael Greenberg for helpful comments on this manuscript.

## Material and Methods

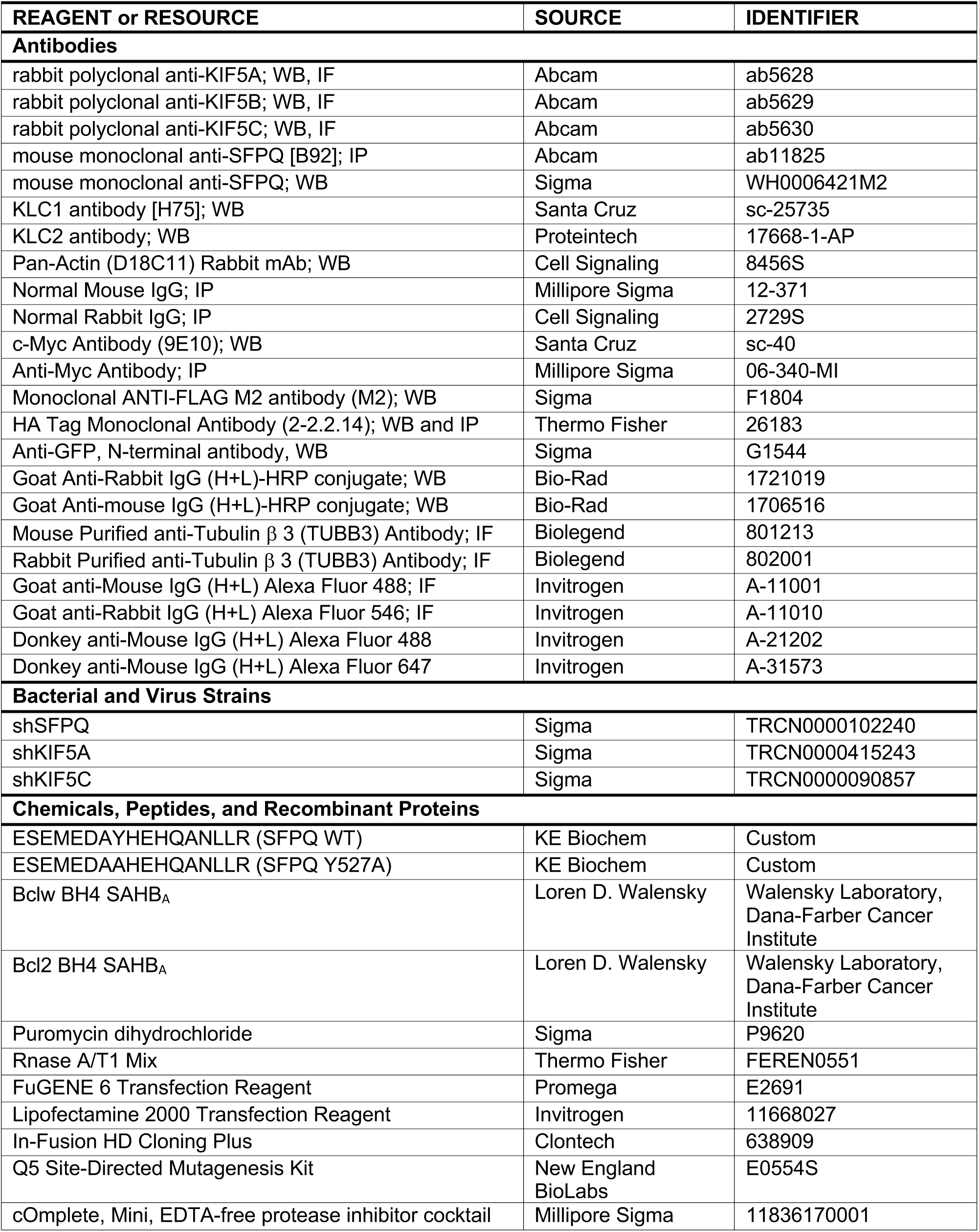
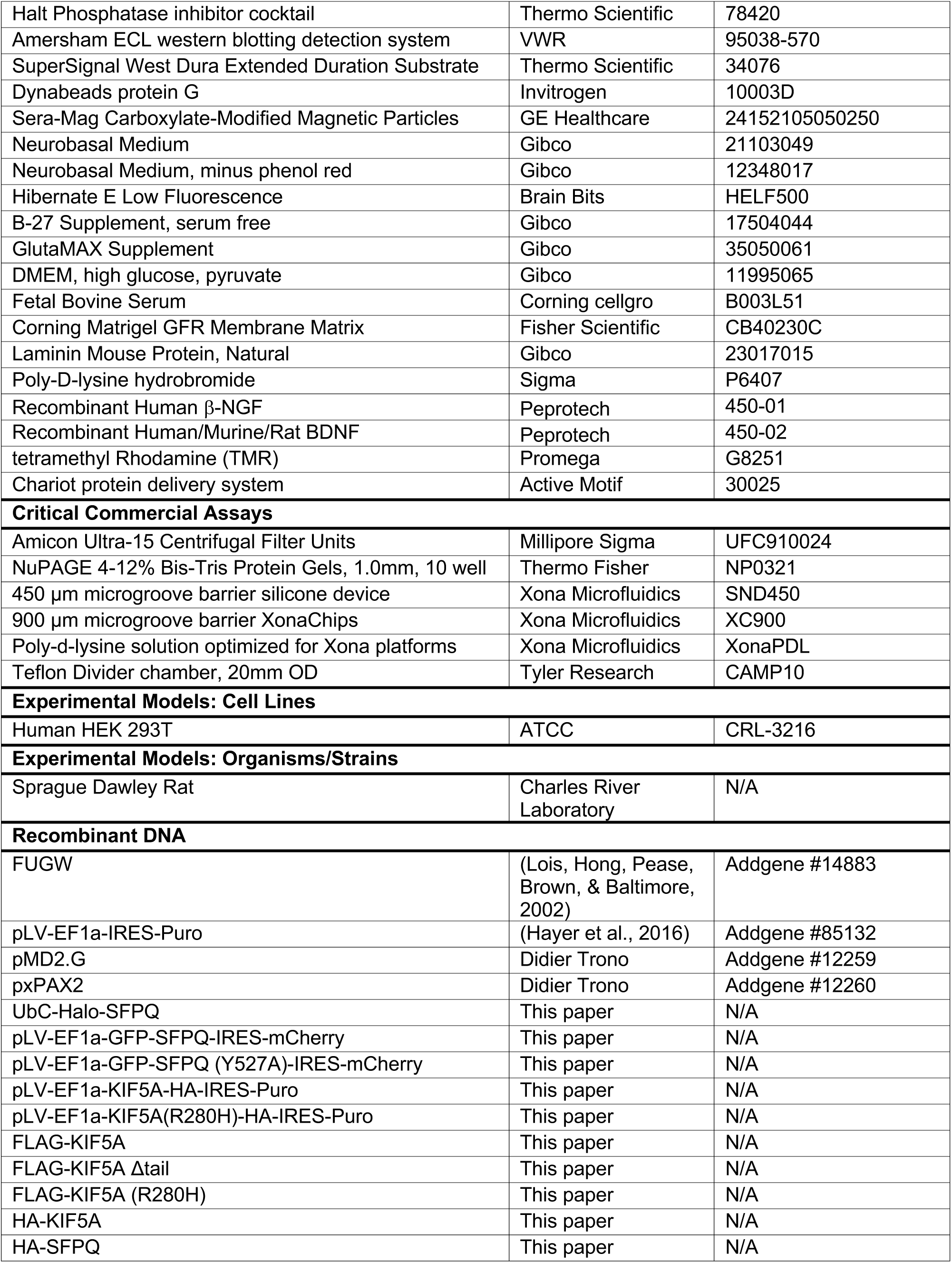
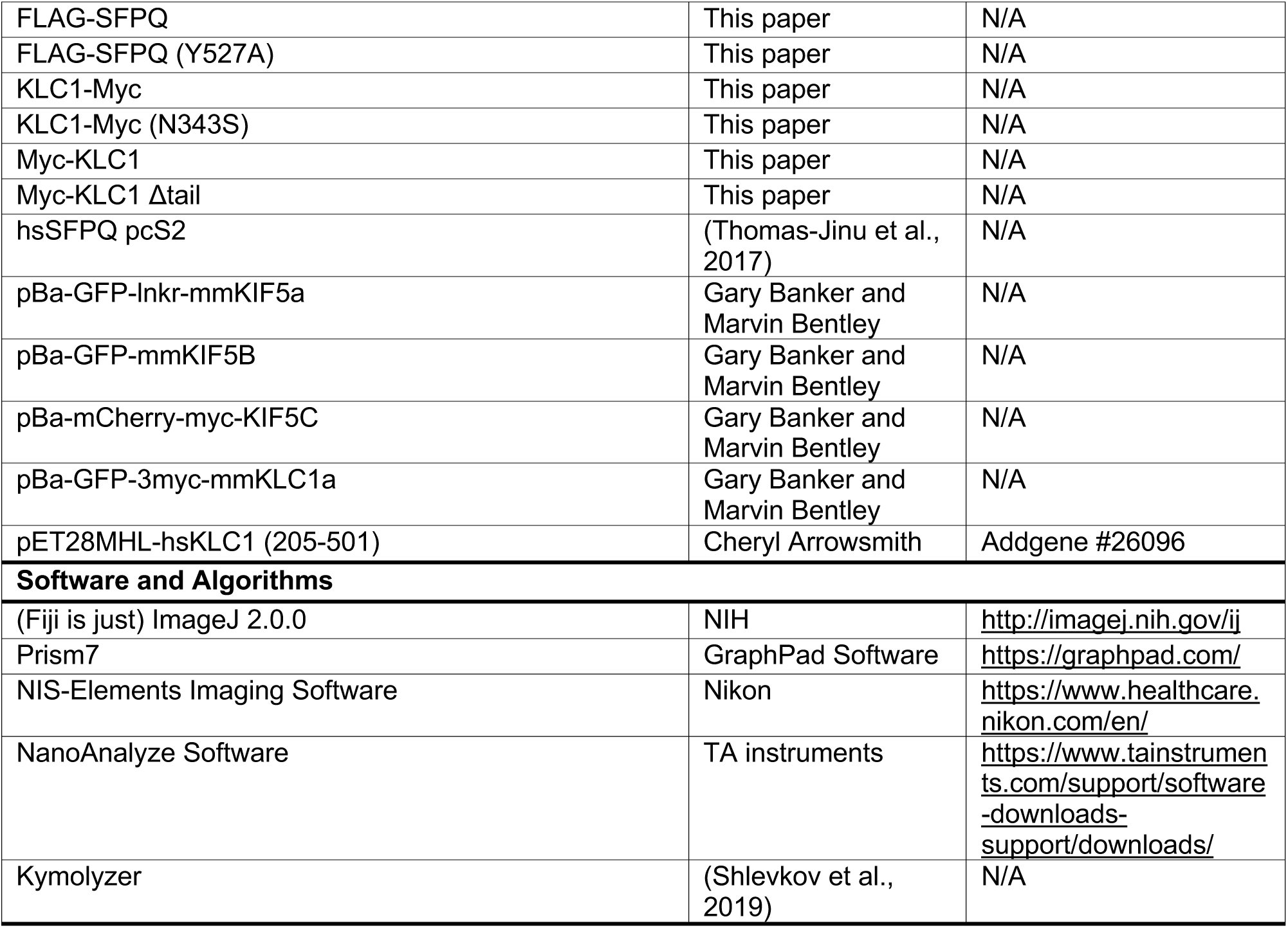
Table for Reagents and Resources.

| REAGENT or RESOURCE | SOURCE | IDENTIFIER |
| --- | --- | --- |
| **Antibodies** |  |  |
| rabbit polyclonal anti-KIF5A; WB, IF | Abcam | ab5628 |
| rabbit polyclonal anti-KIF5B; WB, IF | Abcam | ab5629 |
| rabbit polyclonal anti-KIF5C; WB, IF | Abcam | ab5630 |
| mouse monoclonal anti-SFPQ [B92]; IP | Abcam | ab11825 |
| mouse monoclonal anti-SFPQ; WB | Sigma | WH0006421M2 |
| KLC1 antibody [H75]; WB | Santa Cruz | sc-25735 |
| KLC2 antibody; WB | Proteintech | 17668-1-AP |
| Pan-Actin (D18C11) Rabbit mAb; WB | Cell Signaling | 8456S |
| Normal Mouse IgG; IP | Millipore Sigma | 12-371 |
| Normal Rabbit IgG; IP | Cell Signaling | 2729S |
| c-Myc Antibody (9E10); WB | Santa Cruz | sc-40 |
| Anti-Myc Antibody; IP | Millipore Sigma | 06-340-MI |
| Monoclonal ANTI-FLAG M2 antibody (M2); WB | Sigma | F1804 |
| HA Tag Monoclonal Antibody (2-2.2.14); WB and IP | Thermo Fisher | 26183 |
| Anti-GFP, N-terminal antibody, WB | Sigma | G1544 |
| Goat Anti-Rabbit IgG (H+L)-HRP conjugate; WB | Bio-Rad | 1721019 |
| Goat Anti-mouse IgG (H+L)-HRP conjugate; WB | Bio-Rad | 1706516 |
| Mouse Purified anti-Tubulin $\beta$ 3 (TUBB3) Antibody; IF | Biolegend | 801213 |
| Rabbit Purified anti-Tubulin $\beta$ 3 (TUBB3) Antibody; IF | Biolegend | 802001 |
| Goat anti-Mouse IgG (H+L) Alexa Fluor 488; IF | Invitrogen | A-11001 |
| Goat anti-Rabbit IgG (H+L) Alexa Fluor 546; IF | Invitrogen | A-11010 |
| Donkey anti-Mouse IgG (H+L) Alexa Fluor 488 | Invitrogen | A-21202 |
| Donkey anti-Mouse IgG (H+L) Alexa Fluor 647 | Invitrogen | A-31573 |
| **Bacterial and Virus Strains** |  |  |
| shSFPQ | Sigma | TRCN0000102240 |
| shKIF5A | Sigma | TRCN0000415243 |
| shKIF5C | Sigma | TRCN0000090857 |
| **Chemicals, Peptides, and Recombinant Proteins** |  |  |
| ESEMEDAYHEHQANLLR (SFPQ WT) | KE Biochem | Custom |
| ESEMEDAAHEHQANLLR (SFPQ Y527A) | KE Biochem | Custom |
| Bclw BH4 SAHB<sub>A</sub> | Loren D. Walensky | Walensky Laboratory,<br>Dana-Farber Cancer<br>Institute |
| Bcl2 BH4 SAHB<sub>A</sub> | Loren D. Walensky | Walensky Laboratory,<br>Dana-Farber Cancer<br>Institute |
| Puromycin dihydrochloride | Sigma | P9620 |
| Rnase A/T1 Mix | Thermo Fisher | FEREN0551 |
| FuGENE 6 Transfection Reagent | Promega | E2691 |
| Lipofectamine 2000 Transfection Reagent | Invitrogen | 11668027 |
| In-Fusion HD Cloning Plus | Clontech | 638909 |
| Q5 Site-Directed Mutagenesis Kit | New England<br>BioLabs | E0554S |
| cOmplete, Mini, EDTA-free protease inhibitor cocktail | Millipore Sigma | 11836170001 |
| Halt Phosphatase inhibitor cocktail | Thermo Scientific | 78420 |
| Amersham ECL western blotting detection system | VWR | 95038-570 |
| SuperSignal West Dura Extended Duration Substrate | Thermo Scientific | 34076 |
| Dynabeads protein G | Invitrogen | 10003D |
| Sera-Mag Carboxylate-Modified Magnetic Particles | GE Healthcare | 24152105050250 |
| Neurobasal Medium | Gibco | 21103049 |
| Neurobasal Medium, minus phenol red | Gibco | 12348017 |
| Hibernate E Low Fluorescence | Brain Bits | HELF500 |
| B-27 Supplement, serum free | Gibco | 17504044 |
| GlutaMAX Supplement | Gibco | 35050061 |
| DMEM, high glucose, pyruvate | Gibco | 11995065 |
| Fetal Bovine Serum | Corning cellgro | B003L51 |
| Corning Matrigel GFR Membrane Matrix | Fisher Scientific | CB40230C |
| Laminin Mouse Protein, Natural | Gibco | 23017015 |
| Poly-D-lysine hydrobromide | Sigma | P6407 |
| Recombinant Human β-NGF | Peprtech | 450-01 |
| Recombinant Human/Murine/Rat BDNF | Peprtech | 450-02 |
| tetramethyl Rhodamine (TMR) | Promega | G8251 |
| Chariot protein delivery system | Active Motif | 30025 |
| **Critical Commercial Assays** |  |  |
| Amicon Ultra-15 Centrifugal Filter Units | Millipore Sigma | UFC910024 |
| NuPAGE 4-12% Bis-Tris Protein Gels, 1.0mm, 10 well | Thermo Fisher | NP0321 |
| 450 μm microgroove barrier silicone device | Xona Microfluidics | SND450 |
| 900 μm microgroove barrier XonaChips | Xona Microfluidics | XC900 |
| Poly-d-lysine solution optimized for Xona platforms | Xona Microfluidics | XonaPDL |
| Teflon Divider chamber, 20mm OD | Tyler Research | CAMP10 |
| **Experimental Models: Cell Lines** |  |  |
| Human HEK 293T | ATCC | CRL-3216 |
| **Experimental Models: Organisms/Strains** |  |  |
| Sprague Dawley Rat | Charles River Laboratory | N/A |
| **Recombinant DNA** |  |  |
| FUGW | (Lois, Hong, Pease, Brown, & Baltimore, 2002) | Addgene #14883 |
| pLV-EF1a-IRES-Puro | (Hayer et al., 2016) | Addgene #85132 |
| pMD2.G | Didier Trono | Addgene #12259 |
| pxPAX2 | Didier Trono | Addgene #12260 |
| UbC-Halo-SFPQ | This paper | N/A |
| pLV-EF1a-GFP-SFPQ-IRES-mCherry | This paper | N/A |
| pLV-EF1a-GFP-SFPQ (Y527A)-IRES-mCherry | This paper | N/A |
| pLV-EF1a-KIF5A-HA-IRES-Puro | This paper | N/A |
| pLV-EF1a-KIF5A(R280H)-HA-IRES-Puro | This paper | N/A |
| FLAG-KIF5A | This paper | N/A |
| FLAG-KIF5A Δtail | This paper | N/A |
| FLAG-KIF5A (R280H) | This paper | N/A |
| HA-KIF5A | This paper | N/A |
| HA-SFPQ | This paper | N/A |
| FLAG-SFPQ | This paper | N/A |
| FLAG-SFPQ (Y527A) | This paper | N/A |
| KLC1-Myc | This paper | N/A |
| KLC1-Myc (N343S) | This paper | N/A |
| Myc-KLC1 | This paper | N/A |
| Myc-KLC1 Δtail | This paper | N/A |
| hsSFPQ pcS2 | (Thomas-Jinu et al., 2017) | N/A |
| pBa-GFP-Inkr-mmKIF5a | Gary Banker and Marvin Bentley | N/A |
| pBa-GFP-mmKIF5B | Gary Banker and Marvin Bentley | N/A |
| pBa-mCherry-myc-KIF5C | Gary Banker and Marvin Bentley | N/A |
| pBa-GFP-3myc-mmKLC1a | Gary Banker and Marvin Bentley | N/A |
| pET28MHL-hsKLC1 (205-501) | Cheryl Arrowsmith | Addgene #26096 |
| **Software and Algorithms** |  |  |
| (Fiji is just) ImageJ 2.0.0 | NIH | http://imagej.nih.gov/ij |
| Prism7 | GraphPad Software | https://graphpad.com/ |
| NIS-Elements Imaging Software | Nikon | https://www.healthcare.nikon.com/en/ |
| NanoAnalyze Software | TA instruments | https://www.tainstruments.com/support/software-downloads-support/downloads/ |
| Kymolyzer | (Shlevkov et al., 2019) | N/A |

All experimental procedures were done in accordance with the National Institute of Health guidelines and were approved by the Dana-Farber Cancer Institutional Animal Care and Use Committee.

### Animal Use

Time pregnant Sprague-Dawley rats were purchased from Charles River.

### DNA and shRNA constructs

Constructs used for HEK 293T IP studies were cloned into pcDNA3.1 vector using PCR-based In-Fusion HD cloning (Clontech). KIF5 (pBa-GFP-lnkr-mmKIF5A; pBa-GFP-mmKIF5B; pBa-mCherry-myc-KIF5C) and KLC1 (pBa-GFP-3myc-mmKLC1a) constructs were a gift from Dr. Garry Banker and Dr. Marvin Bentley and were used as a template to clone HA, FLAG or Myc-tagged constructs: HA-KIF5A, FLAG-KIF5A (WT, Δtail, and R280H), KLC1-Myc (WT and N343S) and Myc-KLC1 (WT and Δtail). Human version of SFPQ (hsSFPQ pcS2) was a gift from Dr. Corinne Houart and was used to clone HA and FLAG-tagged constructs: HA-SFPQ, and FLAG-SFPQ (WT and Y527A). The R280H KIF5A, N343S KLC1 and Y527A SFPQ mutants were generated by Q5 site-directed mutagenesis (NEB) using manufacturer’s instructions. Lentiviral constructs used for compartmented Campenot cultures were cloned into pLV-EF1a-IRES-Puro, a gift from Tobias Meyer (Addgene plasmid #85132), for KIF5A-HA (WT and R280H) or into pLV-EF1a-IRES-mCherry for GFP-SFPQ (WT and Y527A). For live cell imaging of SFPQ, Halo-tagged SFPQ was cloned into FUGW, a gift from David Baltimore (Addgene plasmid #14883), as a backbone vector. The shRNAs against shSFPQ (TRCN0000102240), KIF5A (TRCN0000415243) and KIF5C (TRCN0000090857) were purchased from Mission.

### DRG sensory neuron culture

DRGs from embryonic day 15 rats of either sex were dissected and trypsinized. For mass cultures, 300,000 cells were plated onto p35 dishes coated with Corning Matrigel GFR Membrane Matrix (Thermo Fisher) diluted in DMEM (Thermo Fisher). Cultures were maintained in Neurobasal (Invitrogen) with 2% B27 supplement (Invitrogen), 1% Penicillin/streptomycin, 1% GlutaMAX (Life Technologies) and 0.08% glucose at 37°C and 7.5% CO_2_. DRG neurons were plated with 0.3 μM AraC; 100 ng/mL NGF (Peptrotech) + BDNF (Peptrotech); and on DIV2 neurotrophins were reduced to 10 ng/mL NGF + BDNF with AraC and maintained until DIV6 for collection and lysis.

For compartmented Campenot chambers, 120,000 cells were plated in the cell body compartment of a Teflon divider (Camp10, Tyler Research) attached to a p35 dish coated with Corning Matrigel GFR Membrane Matrix diluted in DMEM. DRG neurons were initially plated with 0.3 μM AraC; 100 ng/mL NGF + BDNF; and on DIV1 neurotrophins were reduced to 10 ng/mL NGF + BDNF with AraC only in the cell body compartment. On DIV2 cultures were maintained in the same neurotrophin concentration (10 ng/mL for cell body and 100 ng/mL for distal axons) but without AraC and reduced to 0 ng/mL NGF + BDNF in the cell body compartment and 1 ng/mL NGF + BDNF in the distal axon compartment with AraC from DIV5 to DIV8 for collecting protein lysates or for degeneration assay. As efficient knockdown of SFPQ with shRNA takes up to DIV8, the Campenot cultures used for degeneration assay involving shSFPQ were maintained until DIV12 with the following modifications: from DIV5 to DIV8, neurotrophins were maintained at 10 ng/mL NGF + BDNF for both cell body and axons with AraC and then reduced to 1 ng/mL NGF + BDNF for both cell body and axons with AraC until DIV12.

Microfluidic device with 900 μm microgroove barrier XonaChip (XC900; Xona) was prepared for live cell imaging following manufacturer’s instructions except following the PBS washes for XonaPDL (Xona) coating, the device was further incubated with 10 μg/mL of laminin (Life Technologies) for 3 hours at 37°C. The device was then washed with PBS and primed with DRG neuron media until plating. In the cell body compartment, 20,000 DRG neurons (dissected out from embryonic day 14 rats) were plated to attach at room temperature for 5 min. DRG neurons were infected with lentivirus in the cell body compartment diluted in media containing 0.3 μM AraC; 100 ng/mL NGF + BDNF and fresh media with 0.3 μM AraC; 100 ng/mL NGF + BDNF added to the distal axon compartment. At DIV1, virus was removed and replaced with fresh media to both compartments with the cell body neurotrophins reduced to 10 ng/mL NGF + BDNF with AraC and distal axon compartment with 0.3 μM AraC; 100 ng/mL NGF + BDNF. At DIV2, both cell body and distal axon compartment were kept in 0.3 μM AraC; 5 ng/mL NGF + BDNF until live cell imaging at DIV5.

For immunofluorescence in microfluidic chambers, silicon-based 450μm microgroove barrier (SND450; Xona) was prepared following manufacturer’s instructions with modifications. Sterilized cover glass was coated with 0.2 mg/mL poly-D-lysine (Sigma) overnight, then washed with water and dried. Microfluidic chambers were cleaned by briefly soaking it in ethanol then dried; attached onto PDL-coated glass slide; and the chambers and the wells were filled with 10 ug/mL laminin and incubated for 3 hours at 37°C. The chambers and wells were washed three times with neurobasal media and 30,000 DRG neurons were plated in the cell body compartment with 0.3 μM AraC; 50 ng/mL NGF + BDNF and 0.3 μM AraC; 100 ng/mL NGF + BDNF in the distal axon compartment. At DIV1, both compartments were replaced with fresh media with 0.3 μM AraC; 10 ng/mL NGF + BDNF in the cell body compartment and 0.3 μM AraC; 100 ng/mL NGF + BDNF in the distal axon compartment and maintained until DIV5 for immunostaining. See Fenstermacher *et al*. for more details on compartmented culture system (Fenstermacher, Pazyra-Murphy, & Segal, 2015).

### Axonal degeneration assay

Compartmented chamber cultures were fixed at room temperature with 4% PFA diluted 1:2 in DRG neuron media for 10 min, then an additional 20 minutes in undiluted 4% PFA. DRGs were permeabilized with 0.1% Triton X-100 for 10 min; blocked in 3% BSA in 0.1% Triton X-100 for 1 hour at room temperature; and incubated with rabbit anti-Tuj1 (1:400; Biolegend) overnight at 4°C. Cultures were then incubated with secondary antibodies (1:1000; Invitrogen) for 1 hour at room temperature and stained briefly with DAPI. Images of distal axon tips were obtained using a 40X air objective, and axonal degeneration was quantified as a degeneration index, as previously described (Cosker et al., 2016; Sasaki, Vohra, Lund, & Milbrandt, 2009). For peptide rescue experiment, the following peptides synthesized as previously described (Barclay et al., 2015; Pease-Raissi et al., 2017) were used:

Bclw BH4 SAHB_A_ ALVADFVGYKLRXKGYXBGA Bcl2 BH4 SAHB_A_ EIVBKYIHYKLSXRGYXWDA (differential placement of all-hydrocarbon staples (X) along the BH4 sequences of Bclw and Bcl2; and B represents norleucine, replacing the native cysteine and methionine on Bclw and Bcl2, respectively)

The peptides were introduced into axons using 2 μL Chariot protein transfection system (Active Motif) only in the distal axon compartment of compartmented Campenot chambers immediately after R280H virus removal at DIV2. The culture was kept with the peptides until DIV8 for axon degeneration assay.

### Immunofluorescence

DRG neurons grown in silicon-based microfluidic chambers were fixed with 4% PFA in PBS for 20 minutes at room temperature then washed three times with PBS. The device was then carefully removed and then cultures were immediately permeabilized with 0.1% triton in PBS for 10 minutes at room temperature and blocked for 1 hour at room temperature with 3% BSA in PBS. KIF5 primary antibodies (1:100; abcam) and mouse TUJ1 (1:400; Biolegend) diluted in 3% BSA in PBS were incubated at 4°C overnight. The slides were washed three times with PBS and secondary antibodies diluted in 3% BSA in PBS (1:1000; Invitrogen) were incubated for 1 hour at room temperature. Finally, the slides were washed two times with PBS and incubated with DAPI (1:1000) in PBS; washed briefly and mounted. Images were acquired with Nikon C2 S*i* laser-scanning confocal microscope with 60x oil objective using NIS-Elements imaging software.

### Whole mount immunostaining

Whole DRG with peripheral nerves were dissected from P1 mice of either sex and fixed with 4% PFA at 4°C overnight. DRGs were washed in PBS, permeabilized in 0.5% Triton X-100 for 1 hour and blocked in 5% BSA and 0.5% Triton X-100 for 4 hours. DRGs were incubated for 48 hours in primary KIF5 (1:50, Abcam) and TUJ1 (1:300, Biolegend) antibodies at 4°C and washed overnight in PBS. DRGs were then incubated in secondary antibody (1:500; Life Technologies) at room temperature for 3.5 hours. Images were acquired with Nikon C2 S*i* laser-scanning confocal microscope with 40x oil objective.

### Live cell imaging

DRGs neurons infected with lentivirus expressing Halo-SFPQ were grown in XonaChip microfluidic chambers and were labeled with tetramethyl Rhodamine (TMR) Halo Tag ligand (Promega) according to the manufacturer’s instructions with modifications. TMR stock was diluted in DRG culture media at 1:200 and used at a final labeling concentration of 2.5 μM added to both cell body and axon compartment. DRGs were incubated for 15 mins at 37°C and washed 3 times with complete culture media made with neurobasal without phenol red and incubated for 30 mins at 37°C to wash unbound ligand. DRG culture media was replaced with low fluorescence imaging media (HibernateE; Brain Bits) supplemented with 2% B27 and 1% GlutaMAX. DRG neurons were imaged live in an environmental chamber at 37°C and 7.5% CO_2_ using a 60x oil 1.4NA objective with a Perfect Focus System one frame every 1.5 sec for 3 mins. Images were analyzed using Kymolyzer macro for ImageJ developed by the laboratory of Dr. Thomas Schwarz (Shlevkov et al., 2019). To determine the SFPQ granule diameter, measurement was taken from the two edges of the granule running parallel to the direction of the axon.

### Western blot

HEK 293T cells or DRG sensory neurons were collected and prepped with lysis buffer (1% NP-40; 50mM Tris-HCl, pH 7.4; 150mM NaCl; 2mM EDTA; protease inhibitor (Sigma); and phosphatase inhibitor (Life Technologies)). Cell lysates were placed on ice for 20 minutes and centrifuged at 13,000 rpm for 20 minutes to collect the supernatant. The lysates were separated by 4-12% Bis-Tris NuPAGE gel (Thermo Fisher) and blotted with the following primary antibodies: mouse anti-SFPQ (1:1000; Sigma), rabbit anti-KIF5A (1:2000; Abcam), rabbit anti-KIF5B (1:2000; Abcam), rabbit anti-KIF5C (1:2000; Abcam), rabbit anti-KLC1 (1:500; Santa Cruz), rabbit anti-KLC2 (1:1000; Proteintech), rabbit anti-pan actin (1:1000; Cell Signaling), mouse anti-Myc (1:500; Santa Cruz), mouse anti-FLAG (1:1000; Sigma), mouse anti-HA (1:10,000; Thermo Fisher), rabbit anti-GFP (1:2000; Sigma). HRP-conjugated secondary antibodies (1:10,000; BioRad); ECL detection system (VWR) and SuperSignal West Dura (Thermo Fisher) were used for chemiluminescent detection.

### Transfection and immunoprecipitation

HEK 293T cells were cultured in a 10 cm plate with DMEM, 10% FBS (Thomas Scientific) and 1% Penicillin/streptomycin at 37°C and 5% CO_2_. For transfection, cells were plated in a 6 cm dish and 24 hours later plasmids were transfected with Lipofectamine 2000 (Invitrogen) based on manufacture’s protocol and then incubated for 24 hours before immunoprecipitation experiments.

For immunoprecipitation, HEK 293T cell or DRG neurons were collected and lysed as described for western blots, and 500 μg of protein lysate was precleared with 3 μL of Dynabeads protein G (Thermo Fisher) for 1 (HEK 293T lysates) or 2 hours (DRG neuron lysates) nutated at 4°C. For RNase experiments, the lysate was treated with RNase A/T1 (Fisher Scientific) for 1 hour at room temperature and immediately immunoprecipitated. Following the manufacturer’s instructions, protein lysate was immunoprecipitated for 2 hours at 4°C with the following antibodies: rabbit anti-myc (2.5 μg, Millipore Sigma), mouse anti-HA (2.5 μg, Thermo Fisher), mouse anti-SFPQ (20 μg, Abcam), control normal mouse (Millipore Sigma) or normal rabbit IgG (Cell Signaling),. The input (0.5% for HEK 293T and 3% for DRG) and the elute was analyzed by western blot.

### Lentivirus production and infection

HEK 293T cells grown on 10 cm dish were transfected using FuGENE 6 (Promega) with the transfer vector pxPAX2 (Addgene #12260) and pMD2.G (Addgene plasmid #12259), gifts from Didier Trono, at a ratio of 4:3:1. The transfection reagent was replaced with fresh media after 24 hours. Virus-containing media were collected 48- and 72-hours post transfection; pooled; centrifuged at 1200 rpm for 5 minutes; and filtered through a 0.45 μm PES filter. Finally, the virus was concentrated using Amicon ultra-15 centrifugal filter units (Millipore Sigma) by centrifuging at 3000 rpm and stored at -80°C until use. For infection of DRG sensory neurons, virus was added at DIV1 for 24 hours, except for XonaChip where virus was added immediately after plating. For experiments involving puromycin (Sigma) selection (Figure 6D, shSFPQ; Figure 6F, KIF5A WT and R280H constructs; Figure 2-figure supplement 1C and 1D, shKIF5A and shKIF5C; Figure 6-figure supplement 1, shSFPQ), the neurons were allowed to recover for 1 day after virus removal and 0.4 μg/mL puromycin was then added at DIV3 and replaced with fresh media at DIV5.

### Protein expression and purification

A construct of human KNS2 covering residues 205-501 in the pET28MHL vector, a gift from Cheryl Arrowsmith (Addgene plasmid #26096) was expressed in E. coli BL21 (DE3) in TB medium in the presence of 50 μg/mL of kanamycin. Cells were grown at 37°C to an OD of 0.6, induced overnight at 17°C with 400 μM isopropyl-1-thio-D-galactopyranoside, collected by centrifugation, and stored at -80°C. Cell pellets were microfluidized at 15,000 psi in buffer A1 (25mM HEPES (7.5), 500mM NaCl, 5% glycerol, 30mM Imidazole, 5uM ZnAc, and 7mM BME) and the resulting lysate was centrifuged at 13,000 rpm for 40 min. Ni-NTA beads (Qiagen) were mixed with lysate supernatant for 45 min, washed with buffer A1, and eluted with buffer Bi (25mM HEPES (7.5), 500mM NaCl, 5% glycerol, 400mM Imidazole, 5uM ZnAc, and 7mM BME). The sample was gel-filtered through a Superdex-200 16/60 column in buffer A3 (20mM HEPES (7.5), 200mM NaCl, 5% glycerol, 1mM DTT, and 0.5mM TCEP). Fractions were pooled, but protein began precipitating when concentrated. To combat apparent precipitation, excess NaCl solution was added to a final buffer composition of 18mM HEPES-7.5, 680mM NaCl, 4.5% glycerol, 0.9mM DTT, and 0.45mM TCEP. Adjusted sample was then concentrated and stored at -80°C.

### Isothermal Titration Calorimetry (ITC)

All calorimetric experiments were carried out in 20mM HEPES, pH 7.5, 150mM NaCl, and 0.5mM TCEP, with 2% DMSO at 25°C using an Affinity ITC from TA Instruments (New Castle, DE) equipped with auto sampler. Briefly, 350μL of buffer or protein at 20μM was placed into the calorimetric cell, and 250μL of various SFPQ peptides (KE BioChem) at 200μM were loaded into titration syringe. 4μL syringe solution was injected into the calorimetric cell 30 times with a 200 second interval between injections. Thermodynamic parameters (Kd, stoichiometry, and enthalpy) were calculated according to the single site model provided in NanoAnalyze software (TA instruments).

### Mass Spectrometry

Antibody-conjugated protein G beads from KLC1 and SFPQ immunoprecipitates were suspended in 100 µL of ammonium bicarbonate, reduced with 10 mM dithio-treitol (DTT) for 30 minutes at 56°C and alkylated with 20 mM iodoacetamide for 20 minutes at 22°C in the dark. Excess iodoacetamide was quenched by adding 10 mM dithio-treitol (DTT) before diluting the samples to 250 µL with 100 mM ammonium bicarbonate. Immunoprecipitated proteins were digested overnight at 37°C with 4 µg of trypsin. Tryptic peptides were desalted using 500 µg of a 1:1 mixture of hydrophobic and hydrophilic Sera-Mag Carboxylate-modified Speed Beads (GE Healthcare Life Sciences).

Peptides were loaded onto a precolumn (100 µm × 4 cm POROS 10R2, Applied Biosystems) and eluted with an HPLC gradient (NanoAcquity UPLC system, Waters; 1%–40% B in 90 min; A = 0.2 M acetic acid in water, B = 0.2 M acetic acid in acetonitrile). Peptides were resolved on a self-packed analytical column (30 µm × 50 cm Monitor C18, Column Engineering) and introduced in the mass spectrometer (QExactive HF mass spectrometer, ThermoFisher Scientific) equipped with a Digital PicoView electrospray source platform (New Objective)(Ficarro et al., 2009).

The mass spectrometer was programmed to perform a combination of targeted (Parallel Reaction Monitoring, PRM) and data dependent MS/MS scans. To select precursors for the PRM experiments, we first analyzed a small aliquot of digested KLC1 immunoprecipitate and selected the most intense precursor for peptides mapping uniquely to genes (Askenazi, Marto, & Linial, 2010) encoding each protein of interest. In data-dependent mode, the top 5 most abundant ions in each MS scan were subjected to collision induced dissociation (HCD, 27% normalized collision energy) MS/MS (isolation width = 1.5 Da, intensity threshold = 1E5, max injection time: 50 ms). Dynamic exclusion was enabled with an exclusion duration of 30 seconds. PRM scans were scheduled across an 8 minute-period for each of 45 precursors selected as described above (isolation width = 1.6 Da, max injection time: 119 ms). ESI voltage was set to 3.8 kV.

MS spectra were recalibrated using the background ion (Si(CH3)2O)6 at m/z 445.12 +/-0.03 and converted into a Mascot generic file format (.mgf) using multiplierz scripts (Alexander, Ficarro, Adelmant, & Marto, 2017; Askenazi, Parikh, & Marto, 2009; Parikh et al., 2009). Spectra were searched using Mascot (version 2.6) against three appended databases consisting of: i) rat protein sequences (downloaded from UniProt on 04/09/2018); ii) common lab contaminants and iii) a decoy database generated by reversing the sequences from these two databases. Precursor tolerance was set to 20 ppm and product ion tolerance to 25 mmu. Search parameters included trypsin specificity, up to 2 missed cleavages, fixed carbamidomethylation (C, +57 Da) and variable oxidation (M, +16 Da). Spectra matching to peptides from the reverse database were used to calculate a global false discovery rate and were discarded. Data were further processed to remove peptide spectral matches (PSMs) to the forward database with an FDR greater than 1.0%. Protein abundance for KLC1 and KLC2 (Fig. 1G) or KIF5A, KIF5B, KIF5C (Fig. 1H), in the KLC1 immunoprecipitate was calculated by summing the extracted ion chromatogram peak area of the 3 most abundant (Silva, Gorenstein, Li, Vissers, & Geromanos, 2006) gene-unique peptide sequences and averaged across 2 technical replicates. Due to the low absolute abundance of KLC and KIF proteins in the SFPQ immunoprecipitates, we used an MS2-level quantitation approach whereby the extracted ion chromatogram intensity (calculated as the area under the curve) of a set of precursor/fragment ion pairs (manually selected from the targeted MS/MS experiments) in the SFPQ LC-MS/MS analyses were normalized to their intensity in the KLC1 immunoprecipitate, taking into account the average abundance of each KLC (Fig. 1G) or KIF5 (Fig. 1H) protein measured by the top3 quantitation method described above.

### Quantification and Statistical Analysis

Data are expressed as mean ± s.e.m. To assess statistical significance, data were analyzed by unpaired two-tailed Student’s t test. For multiple comparisons, data were analyzed by one-way ANOVA. Significance was placed at p < 0.05. Statistical analysis was done using Microsoft Excel and GraphPad Prism.

---

**Introduction-figure supplement 1.**
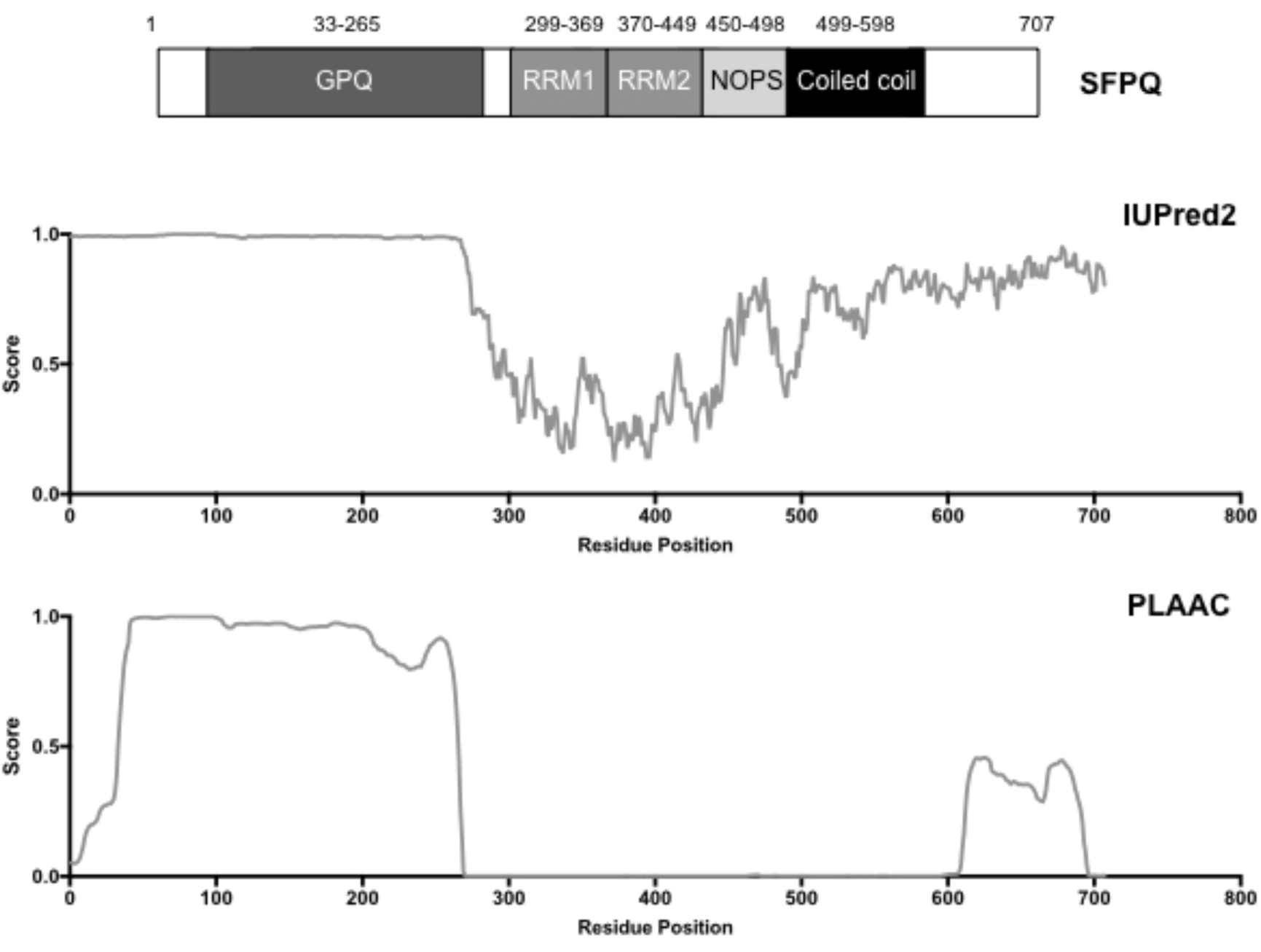
Bioinformatic analysis of SFPQ protein sequence. Schematic of protein domains of SFPQ and the probability score of SFPQ sequence for being disordered and prion-like as predicted by IUPred2 (top) and PLAAC (bottom), respectively.

**Figure 1-figure supplement 1.**
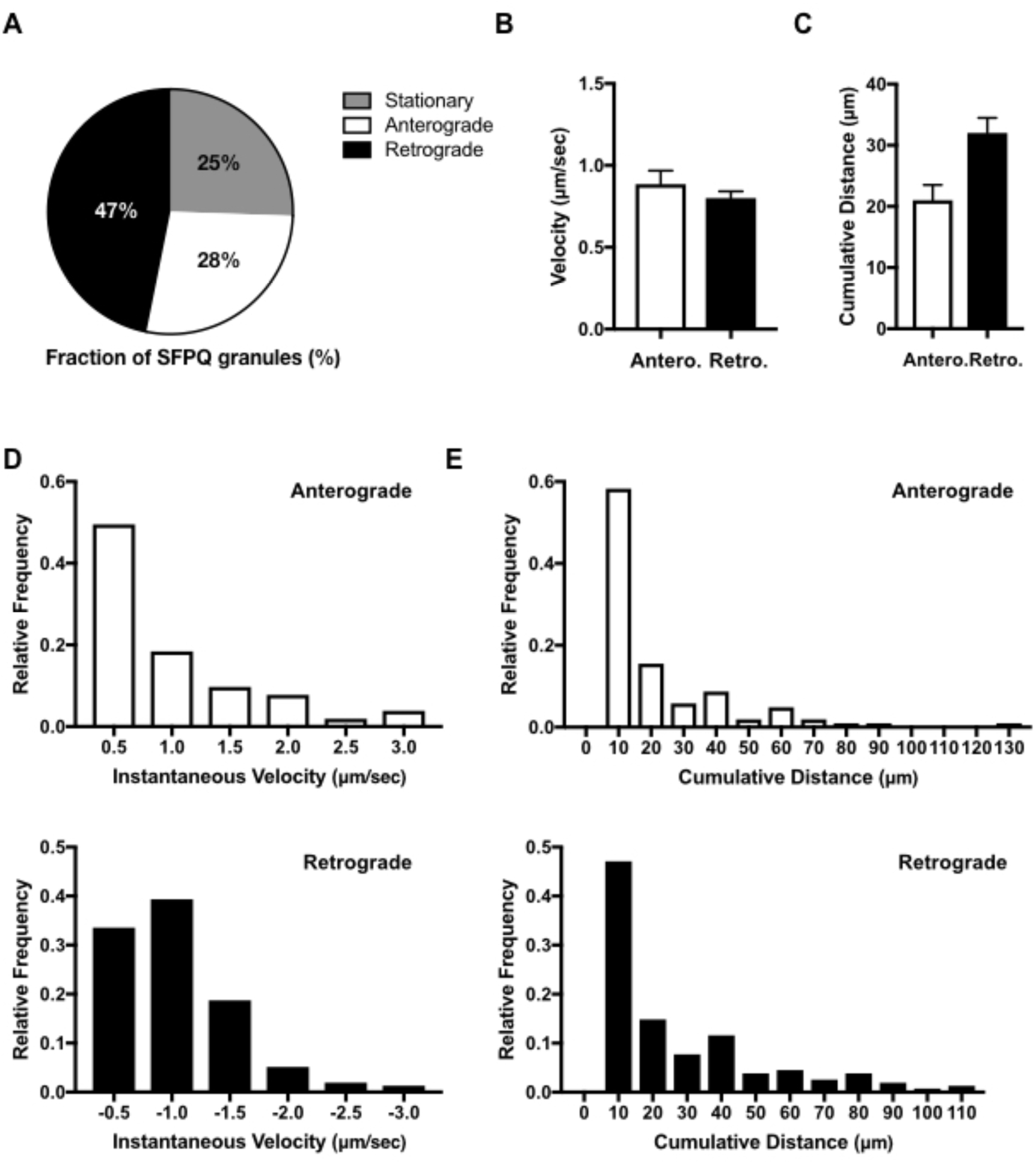
Transport kinetics of SFPQ granules in axons of DRG sensory neurons. (A) Fraction of SFPQ granules spent in stationary, anterograde or retrograde phase. Data analyzed from 217 particles, from 29 axons, across 2 independent experiments. (B) Average velocity and (C) average cumulative distance of Halo-tagged SFPQ granules in axons and its frequency distribution in (D) and (E), respectively.

**Figure 2-figure supplement 1.**
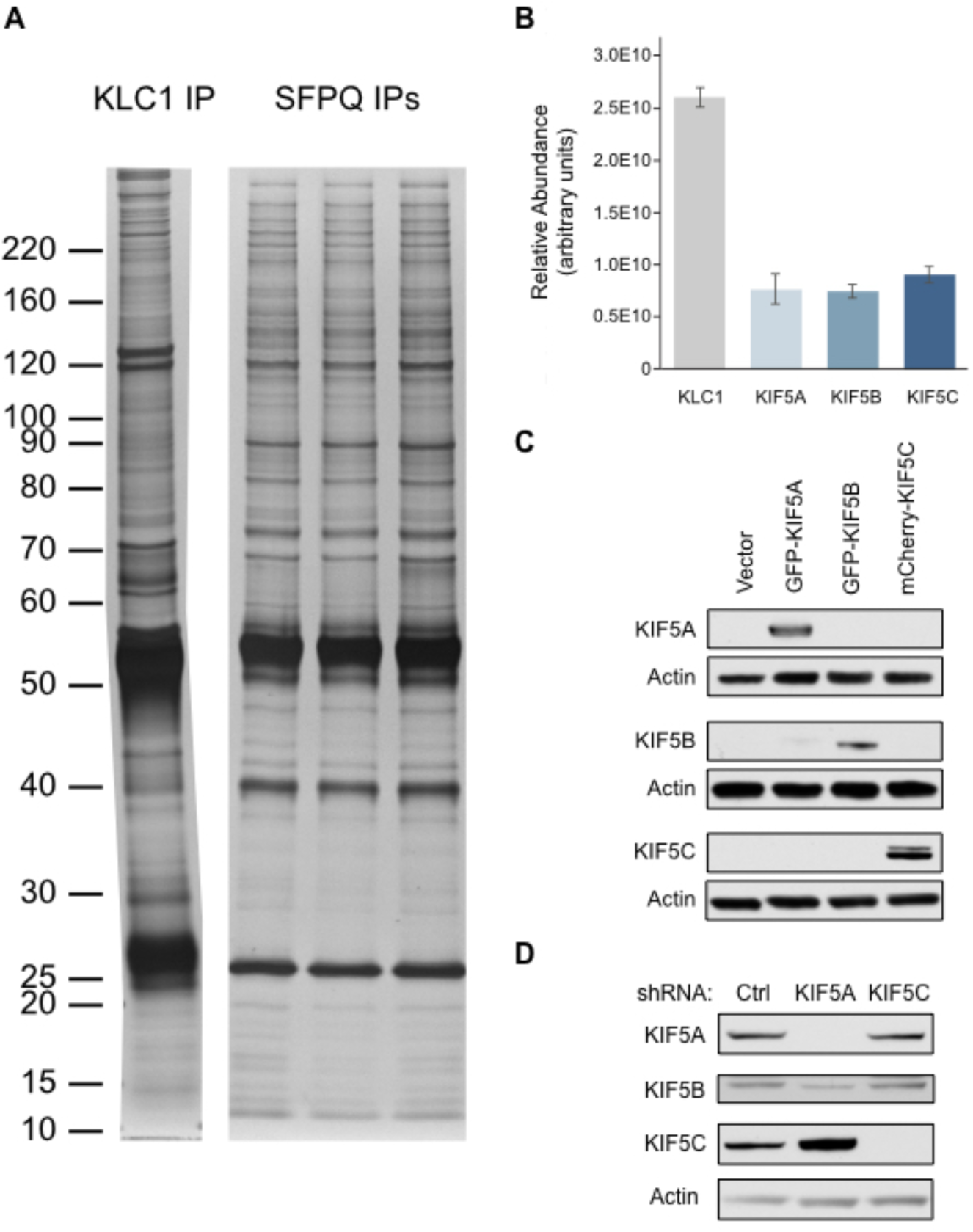
Silver stain analysis of endogenous KLC1 and SFPQ IPs from DRGs and verification of antibodies for KIF5A, KIF5B and KIF5C. (A) Silver stain analysis of endogenous KLC1 and SFPQ proteins purified from DRG extracts. Each lane corresponds to 50% of the enriched material analyzed in Figure 2-figure supplement 1B (KLC1 IP) and Figure 2B and 2C (three independent SFPQ IPs). (B) Relative abundance of KIF5A, KIF5B and KIF5C proteins (with KLC1 as a reference) in a KLC1 IP measured by label-free mass spectrometry. Data represent mean ± s.e.m across two technical replicate analyses. (C) Western blot of HEK 293T lysates transfected with empty vector, GFP-KIF5A, GFP-KIF5B or mCherry-KIF5C and probed with the KIF5 antibodies. Actin serves as loading control. (D) DRG sensory neuron lysates infected with control (Ctrl) or with either shRNA against *KIF5A* or *KIF5C* and probed with the KIF5 antibodies. Actin serves as loading control.

**Figure 2-figure supplement 2.**
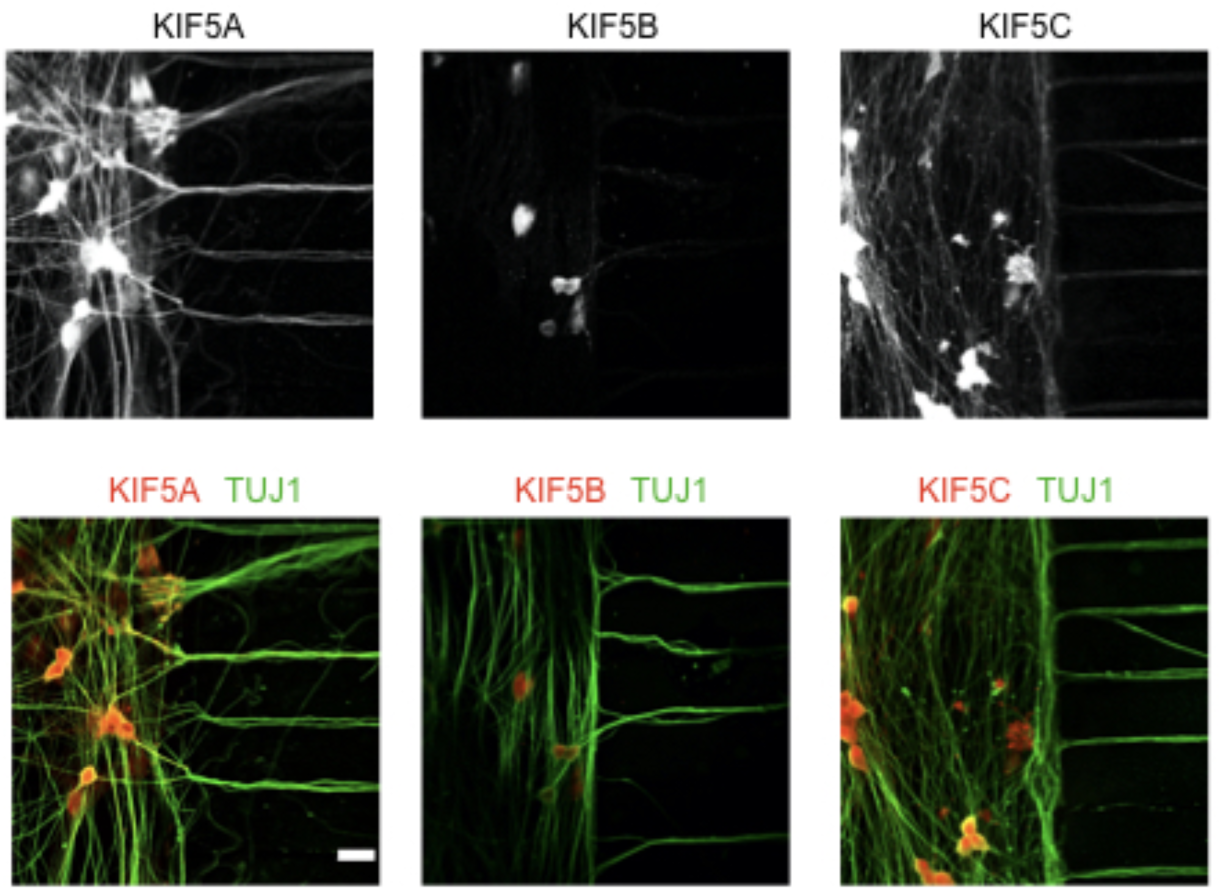
KIF5 motors differentially localize to cell body and distal axons. Representative staining of endogenous KIF5A, KIF5B and KIF5C in DRG sensory neurons grown in microfluidic chambers; Scale bar 20 μm; TUJ1 (green), KIF5 (Red).

**Figure 3-figure supplement 1.**
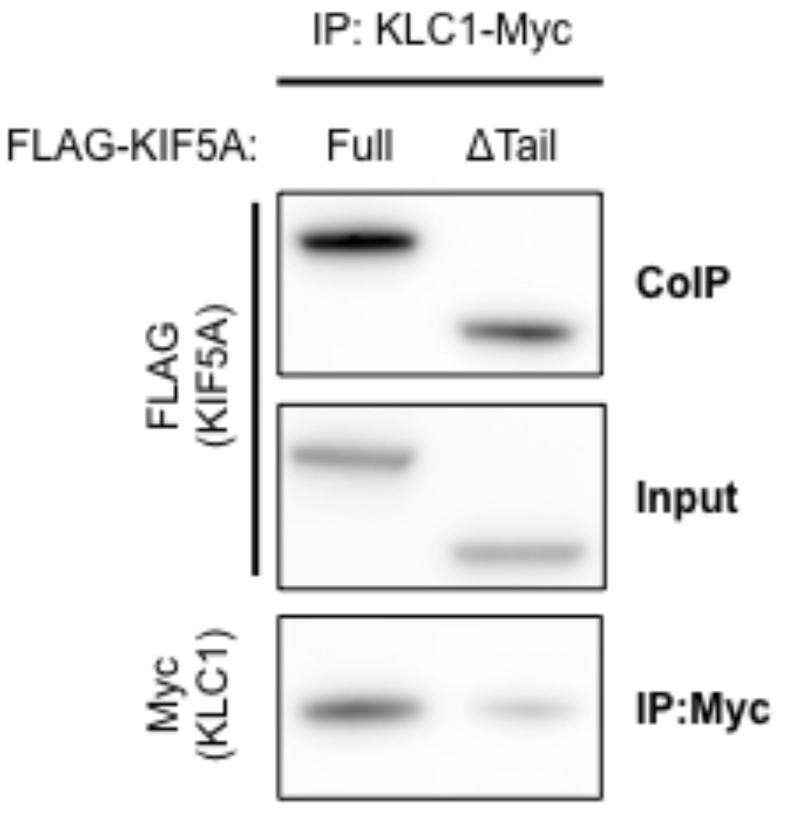
KIF5A ΔTail mutant binds to KLC1. HEK 293T cells transfected with KLC1-Myc and with either WT FLAG-tagged KIF5A or the ΔTail mutant. Myc was IPed and blotted for FLAG and Myc.

**Figure 4-figure supplement 1.**
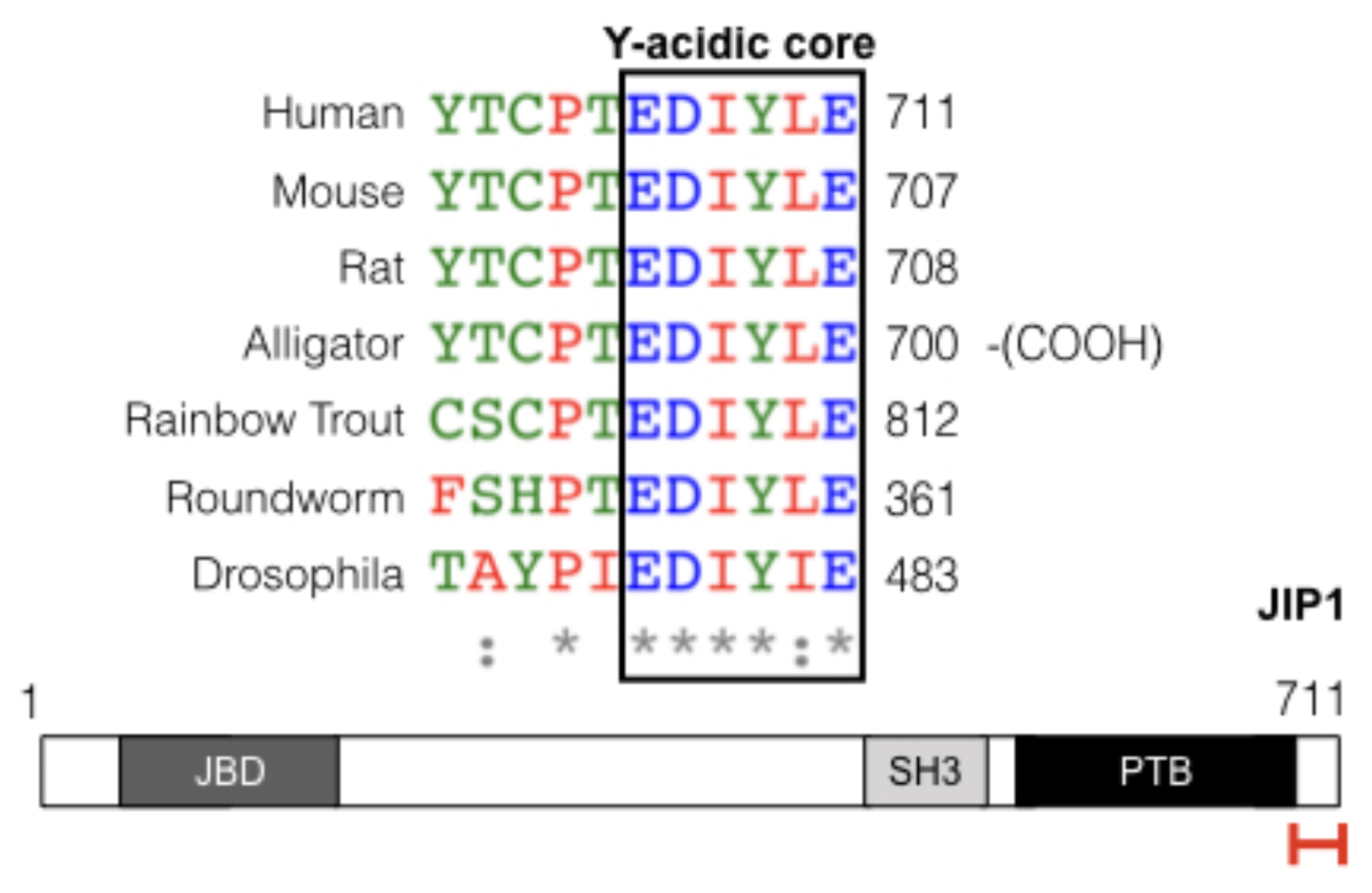
SFPQ and JIP1 both share a Y-acidic motif. Alignment of the sequence within the C-terminal region of JIP1 containing the Y-acidic motif. On the bottom; schematic of the domains of JIP1. Red bracket indicates the region containing the Y-acidic motif. JBD, JNK binding domain; SH3, Src homology-3 domain; PTB, phosphotyrosine binding domain.

**Figure 5-figure supplement 1.**
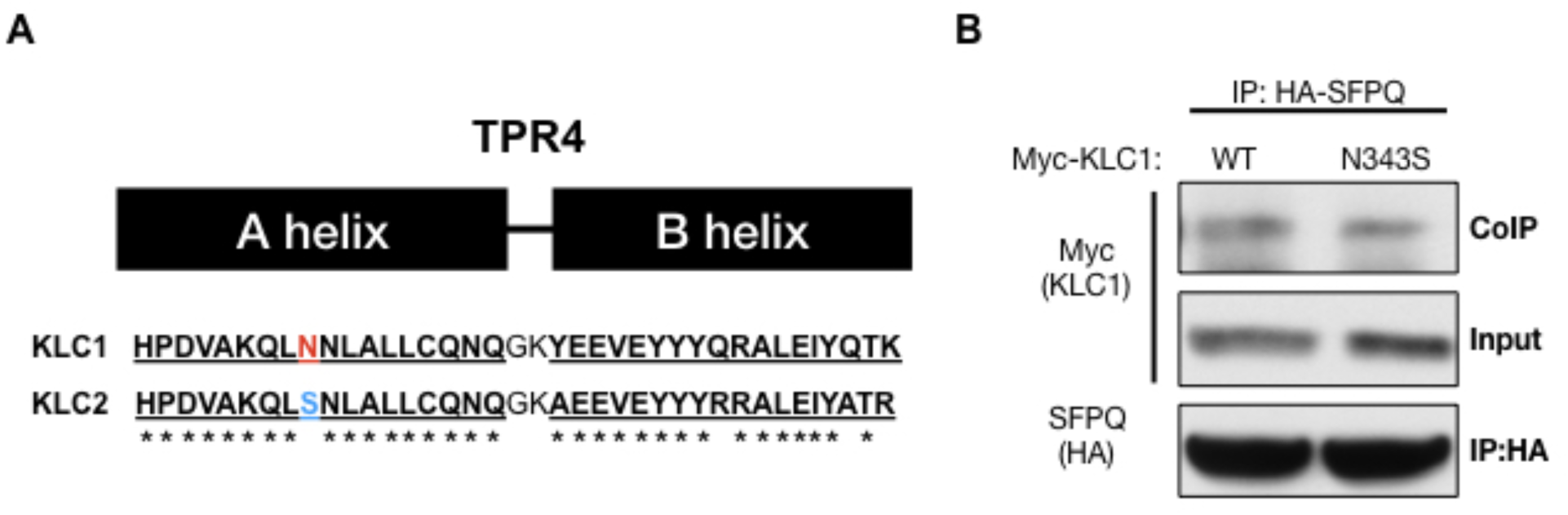
SFPQ mode of binding to KLC1 is distinct from JIP1. (A) Schematic cartoon depicting the location of N343 on KLC1 in comparison to KLC2 on A helix of TPR4. HEK 293T lysates transfected with HA-SFPQ, and with either Myc-tagged WT or N343S mutant of KLC1. HA-SFPQ was IPed and blotted against HA and Myc.

**Figure 6-figure supplement 1.**
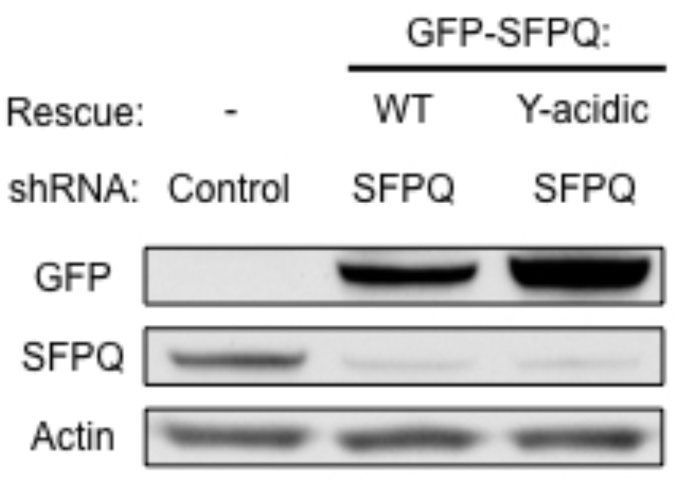
Expression of shRNA-resistant WT or Y-acidic GFP-tagged SFPQ. DRG neurons were infected with control or shRNA against SFPQ and rescued with empty, GFP-tagged WT or Y-acidic mutant of SFPQ. Protein lysates were blotted against GFP, endogenous SFPQ and actin as loading control.

**Figure 6-figure supplement 2.**
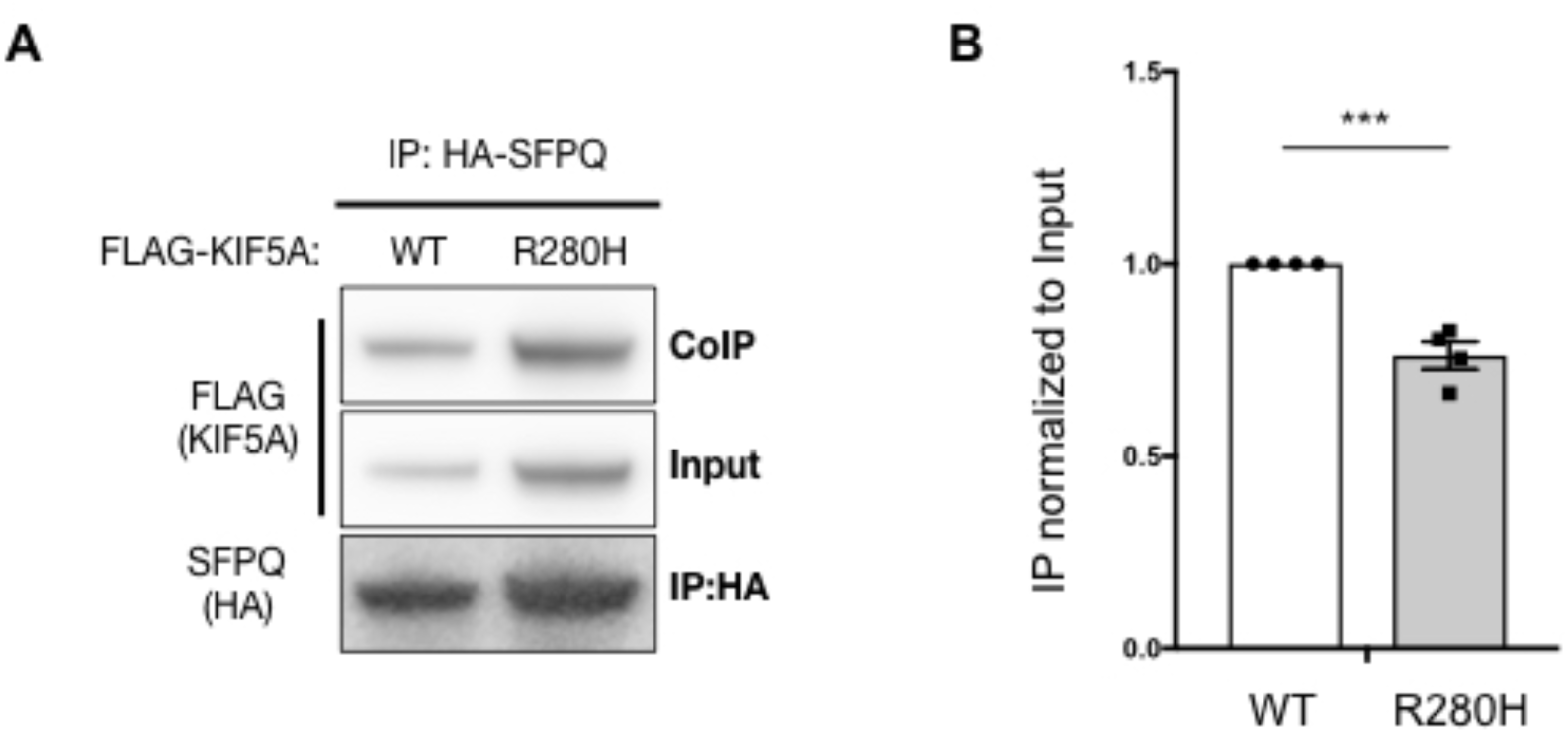
R280H mutation of KIF5A, which impairs transport, also reduces binding to SFPQ. (A) HEK 293T lysates transfected with HA-SFPQ, and with either FLAG-tagged WT or R280H mutant of KIF5A. HA-SFPQ was IPed and blotted against FLAG and HA. (B) Quantification of pull down in (A) normalized to input. ***p = 0.0006 by unpaired two-tailed t test; n = 4; data represent mean ± s.e.m.

**Table1-Source Data 1:** Raw ITC data for WT SFPQ

**Table1-Source Data 2:** Raw ITC data for Y527A Y-acidic mutant

**Video 1:** Halo-SFPQ is transported in anterograde and retrograde manner in axons. Time-lapse movie was captured in axons of dorsal root ganglion sensory neurons grown in compartmented cultures. The video was acquired every 1.5 sec and played at 10 fps.

**Video 2:** Y527A mutation of SFPQ disrupts axonal transport of SFPQ. Time-lapse movie was captured in axons of dorsal root ganglion sensory neurons grown in compartmented cultures. The video was acquired every 1.5 sec and played at 10 fps.

